# A hierarchical activation threshold shapes RSV F antibody repertoires from toddler vulnerability to adult protection

**DOI:** 10.64898/2026.07.28.735825

**Authors:** Jie Deng, Yixin Li, Siyu Lei, Kaijun Xu, Jianhui Nie, Hui Zhai, Jinyue Wang, Lingjie Xu, Fanchong Jian, Rui Feng, Zhe Lv, Wenxiang Yu, Mengyang Ma, Yuxia Zhang, Chuziyue Zhang, Lei Wang, Yuchen Zhang, Yaling Hu, Xiaoyu Xu, Weijin Huang, Enmei Liu, Yunlong Cao, Xiangxi Wang

**Affiliations:** National Laboratory of Biomacromolecules, CAS Center for Excellence in Biomacromolecules, Institute of Biophysics, Chinese Academy of Sciences; Beijing, 100101, China; University of Chinese Academy of Sciences; Beijing, 100049, China; Peking-Tsinghua Center for Life Sciences, Peking University, Beijing, 100871, P.R. China; Biomedical Pioneering Innovation Center (BIOPIC), School of Life Sciences, Peking University, Beijing, 100871, China; Division of HIV/AIDS and Sexually Transmitted Virus Vaccines, National Institutes for Food and Drug Control (NIFDC), No. 31 Huatuo Street, Daxing District, 102629 Beijing, China; Department of Respiratory Medicine, Children’s Hospital of Chongqing Medical University, National Clinical Research Center for Children and Adolescents’ Health and Diseases, Ministry of Education Key Laboratory of Child Development and Disorders; Chongqing, 400014, China; Vazyme Biotech Co., Ltd, Nanjing, 210046, PR China; IBP-Sinovac Joint Laboratory of Cutting-edge Technologies and Vaccine & Drug Development, Beijing, 100101, China; Changping Laboratory, Beijing, 102206, P.R. China; Chongqing Key Laboratory of Child Rare Diseases in Infection and Immunity; Chongqing, 400014, China

## Abstract

Respiratory syncytial virus (RSV) causes severe infant morbidity, yet the mechanisms underlying their suboptimal immunity and the failure of prefusion F-stabilized adult vaccines in infants remain unclear. Here we integrate deep mutational scanning of 761 preF-binding antibodies, repertoire profiling of 102 repeatedly exposed pediatricians and 61 RSV-experienced toddlers, structural analysis, and >40,000 viral genomes to decode RSV F immunity. We resolved 12 immunologically distinct functional antibody subclasses and uncovered a hierarchical activation threshold: apical epitope-targeting antibodies with superior neutralizing potency demand extensive somatic hypermutation and cooperative CDR networks, whereas central-to-basal epitope-directed, marginal or non-neutralizing antibodies engage germline-encoded antibodies through minimal mutations (e.g., S31G). Toddler repertoires are confined to low-threshold, non-productive sites; adult repertoires enrich for apical elite neutralizers whose immune pressure drives contemporary RSV-B evolution. These reframe pediatric RSV vulnerability as a threshold-gated repertoire deficit and prescribe vaccine strategies that actively redirect immunodominance from permissive epitopes toward high-barrier apical targets.

## Introduction

Respiratory syncytial virus (RSV) is a leading cause of severe acute lower respiratory tract infections in infants and older adults worldwide, imposing a substantial and persistent global health burden^1,2^. The viral fusion (F) glycoprotein, which undergoes a conformational change from a metastable prefusion (preF) to a stable postfusion (postF) state, is the primary target for neutralizing antibodies and the focal point of vaccine design^3^. While structural studies have delineated major antigenic sites (Ø–VI) on preF^4,5^, a purely structure-based classification cannot explain the vast functional heterogeneity of antibodies within a single site or predict their protective efficacy. Critically, the mechanistic basis for why naturally infected infants remain susceptible to severe disease, and how repeated viral exposure in adults confers superior protection, has remained elusive. This knowledge gap underpins the historical failures of pediatric RSV vaccines^6-8^ and the persistent unmet need for effective early-life immunization.

The development of preF-stabilized vaccines and potent prophylactic monoclonal antibodies (mAbs) like nirsevimab (site Ø) and clesrovimab (site IV) represents a breakthrough^9-11^. However, a critical disconnect persists: the same preF antigen that successfully boosts protection in seropositive adults has failed to demonstrate sufficient efficacy in seronegative infants, with some candidates even associated with vaccine-enhanced respiratory disease (VAERD)^8,12-16^.This clinical dichotomy mirrors the natural history where primary infection in infants often causes severe disease, while reinfection in adults is typically mild^2,17^. It suggests that the quality of the humoral response is fundamentally shaped by immune maturity and antigen exposure history—factors poorly understood at a functional and repertoire level. Furthermore, while prophylactic antibody drugs are effective, their long-term utility is threatened by potential viral escape^9,18-23^, yet the population-level drivers of RSV antigenic evolution are unknown.

Decoding protective immunity may require understanding the most robust human response. We proposed that pediatric healthcare workers with long-term, high-frequency occupational exposure to RSV represent a model of an optimally matured immune state, enriched for high-potency antibodies shaped by repeated antigenic challenge^24^. Systematically comparing this cohort with immunologically vulnerable toddlers who mount primary, suboptimal responses provides a powerful paradigm to define the hallmarks of a protective repertoire. To move beyond descriptive immunology, we established a high-resolution, functional antigenic landscape of RSV F. We integrated comparative antibody repertoire profiling of these cohorts with deep mutational scanning (DMS)^25,26^, structural analysis, and longitudinal viral genomics. This system-level approach allowed us to define a functional epitope hierarchy based on immunogenicity and somatic hypermutation (SHM) requirements; dissected the divergent immunodominance patterns and antibody maturation pathways between infants and adults; and quantified the immune selection pressures each cohort exerts on circulating RSV strains. These explain the deficits in infant antibody repertoires, delineate the maturation pathways toward broad, potent neutralization in adults, and identify adult-driven immune pressure as a dominant force in contemporary RSV evolution. By linking antibody genetics, epitope function, and host immunobiology to viral escape, this work shifts the paradigm from merely inducing preF-specific responses to strategically eliciting antibodies with the right specificity, potency, and maturity—offering a framework for designing next-generation pediatric vaccines that bridge the infant-adult immunity gap.

## RESULTS

### Toddlers mount preF-specific yet subpotent antibody responses

Primary RSV infection in infants frequently manifests as severe lower respiratory disease, whereas repeated exposure in adults typically yields mild reinfection^2,17^, a dichotomy mirrored by the divergent performance of preF vaccines across age groups^8,12-16^. To define how exposure history shapes humoral quality, we profiled 102 pediatricians with long-term occupational RSV exposure and 61 toddlers (0–3 years old) with confirmed prior RSV infection (Figure 1A). Serum neutralization showed pediatricians exhibited 3–8-fold higher GMTs against four RSV strains versus toddlers (Figure 1B). We isolated RSV F-specific IgG⁺ memory B cells (CD19⁺IgD⁻IgM⁻CD27⁺) by single-cell sorting on preF and postF probes, screened supernatants by ELISA, and sequenced positive wells (Figure S1A). In total, 1,597 variable heavy (VH) and light (VL) pairs from pediatricians and 348 from toddlers were cloned and expressed as full-length IgG. SHM levels increased with age and exposure and pediatrician-derived antibodies showed higher SHM than those from naturally infected adults^27^, indicating that occupational exposure further elevates antibody maturation (Figure S1B).

**Figure 1.**
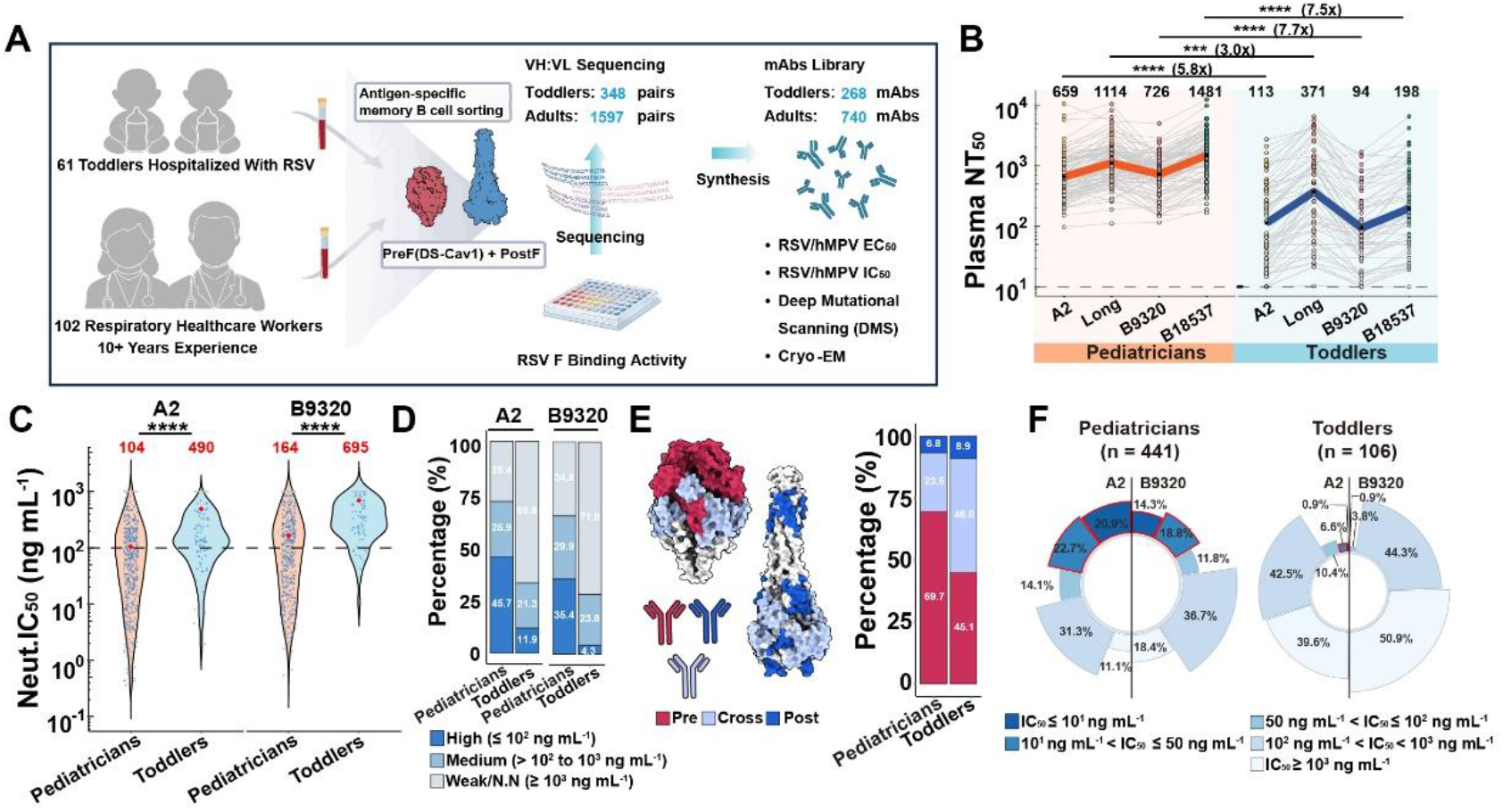
Age-dependent differences in RSV-neutralizing antibody function and binding preference. **(A)** Schematic diagram of antibody isolation and functional characterization. **(B)** Plasma half-maximal neutralizing titers (NT_50_) against four RSV strains (A2, Long, B9320, B18537) in pediatricians and toddlers. Numbers above dots and the connecting line indicate geometric mean NT_50_. **(C)** Neutralization potency (IC_50_) of antibodies from pediatricians and toddlers against RSV A2 and B9320. Red dots and values above indicate the geometric mean IC_50_, and the black dashed line marks the threshold for high neutralizing potency. **(D)** Proportions of neutralizing antibodies against RSV A2 and B9320 in pediatricians and toddlers, grouped by neutralizing potency. N.N., non-neutralizing. **(E)** Schematic diagram of preF-specific, cross-reactive and postF-specific antibody epitopes (left). Proportions of each antibody type in pediatricians and toddlers (right). **(F)** Proportions of preF-specific antibodies against RSV A2 and B9320 in pediatricians and toddlers, grouped by neutralizing potency. ***p<0.001, ****p<0.0001.

Neutralization assays confirmed pediatrician-derived mAbs were ∼5-fold more potent against RSV A2 and B9320 than toddler mAbs (Figure 1C). High-potency antibodies (half-maximal inhibitory concentration, IC_50_ ≤ 100 ng mL^-1^) constituted 46% (A2) and 35% (B9320) of the pediatrician repertoire, but only ∼12% and 4% of the toddler repertoire, which was dominated by low-efficacy ones (Figure 1D). Based on preF/postF binding ratios (half-maximal effective concentrations, EC_50_), we categorized mAbs as preF-specific (ratio < 0.01), cross-reactive (0.01–100), or postF-specific (>100) (Figure 1E; Figure S1C). PreF-specific mAbs were 4–6-fold more potent than cross-reactive mAbs, while postF-specific mAbs were non-neutralizing (Figure S1D). This potency correlated with the highest SHM in the preF-specific subset (Figure S1E). Notably, preF-specific antibodies constituted the dominant fraction of high-potency neutralizers in pediatric donors, displaying mean IC_50_ values of 57 ng mL^-1^ and 101 ng mL^-1^ against RSV A2 and B9320, respectively, and representing ∼70% of the total antibody repertoire (Figure 1E; Figure S1F). Although preF-specific antibodies accounted for nearly half of the toddler antibody repertoire, their neutralization potency was much lower than that of pediatrician-derived equivalents, with mean IC_50_ values of 310 ng mL^-1^ and 560 ng mL^-1^ against RSV A2 and B9320, respectively (Figure 1E; Figure S1F). Strikingly, the difference in extremely potent neutralizer frequency was even more pronounced: 21% (14%) and 44% (33%) of pediatrician-derived antibodies reached IC_50_ ≤ 10 ng mL^-1^ and ≤ 50 ng mL^-1^ against RSV A2 (B9320), compared with 1% (0%) and 8% (1%) of toddler-derived antibodies meeting the same thresholds (Figure 1F). In contrast, cross-reactive antibodies from both cohorts showed no significant differences in neutralization activity and proportional distributions (Figures S1F and S1G). These results demonstrate that toddlers can mount preF-specific responses, but the repertoire is qualitatively distinct, with reduced potency and a scarcity of elite neutralizers, reflecting divergent epitope recognition and maturation trajectories.

### A functional epitope landscape resolves 12 subclasses and an apical-to-basal potency hierarchy

Current antigenic sites of the RSV preF, derived from crystallography and cryo-EM, delineate seven major sites (Ø–VI)^5,16,28^. However, these structure-based classifications suffer from spatial ambiguity, inability to define functional immune escape boundaries, and low throughput, failing to resolve key determinants of antibody potency. Clustering of 23 published and 39 new cryo-EM structures herein confirms that many antibodies map contiguously across multiple canonical sites, with quaternary epitopes like that of AM14^29,30^ defying discrete categorization (Figures S2A and S2B). To overcome these limitations, we performed deep mutational scanning (DMS) on 761 preF-binding antibodies (571 from pediatricians, 172 from toddlers and 18 from public dataset) using yeast (monomer) and mammalian (trimer) display platforms, which showed high concordance (Figure S2C). Graph-based clustering of escape profiles revealed 12 distinct functional subclasses embedded within seven major epitope classes (Ø–VI), expanding the previous classification by five new groups (Figure 2A; Figure S2D). The DMS-defined classes largely recapitulated the spatial organization of structural sites but with critical functional refinements (Figure 2B; Figure S2). Overall, neutralization potency correlates with both epitope location and conformational specificity: apical sites Ø and V show the highest activity; central sites II to IV show intermediate potency (III > II > IV); and basal sites I and VI are largely non-neutralizing (Figure 2B).

**Figure 2.**
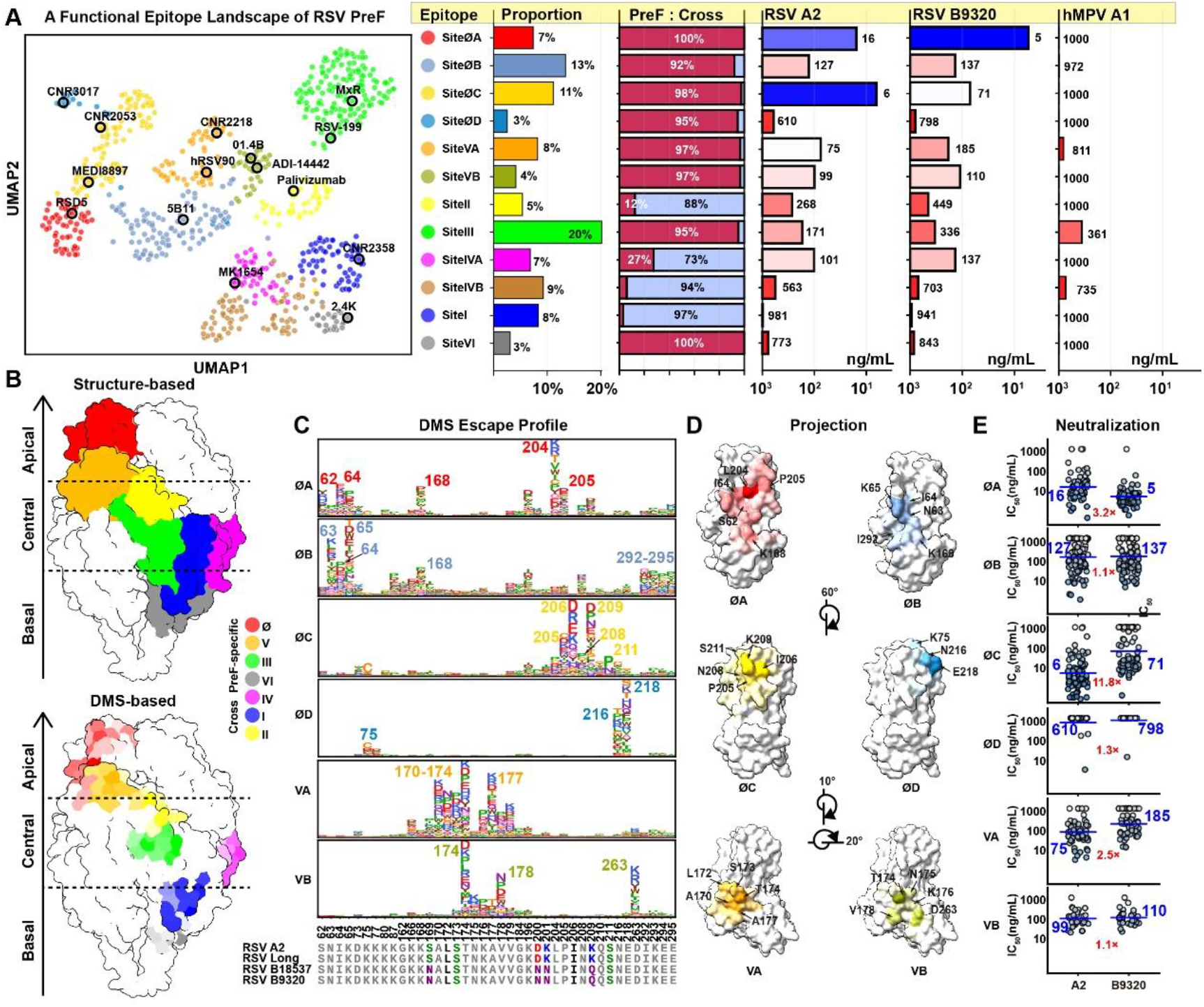
Structural and functional antigenic atlas of RSV F. **(A)** Left: UMAP embedding of epitope groups of monoclonal antibodies binding to RSV preF, including antibodies isolated from our cohorts and previously published preF-binding antibodies (n = 18). Right: Proportion of each epitope cluster, with fractions of preF-specific and cross-reactive antibodies and geometric mean IC_50_ values for each cluster against RSV A2, RSV B9320, and hMPV A1. Bar length represents log_10_ (1000/geometric mean IC_50_), and colors indicate IC_50_ values (blue, low; red, high). **(B)** Projection of classical antigenic sites and newly defined functional epitopes onto the RSV preF structure. Functional epitopes are colored according to their corresponding classical antigenic sites, with color intensity indicating residue-level functional contribution inferred from DMS; darker colors denote greater functional importance. **(C)** Escape mutation profiles of RSV F binding activity derived from deep mutational scanning (DMS) (upper panel) and epitope amino acid multiple sequences alignment analysis for sites Ø and V (lower panel). Escape scores for each epitope are shown in the upper panels, with top escape residues highlighted in epitope-specific colors (Figure 2A). Conserved residues are shown in gray. Variable residues are colored by physicochemical properties: hydrophobic (A, L, I, P, F, M), black; polar (G, S, T), yellow; neutral (Q, N), purple; acidic (D, E), red; basic (R, K), blue. **(D)** Surface mapping of site Ø and V escape risk on RSV F, colored as in **(A)**. Color intensity reflects residue-level DMS escape scores, with darker shades indicating higher escape scores and greater escape risk. Arrows indicate critical escape residues. **(E)** Neutralization potency (IC_50_) of antibodies targeting site Ø and V against RSV A2 and B9320. Blue lines and values indicate the geometric mean IC_50_ of antibodies.

Site Ø comprises four functionally distinct subclasses (ØA–ØD) with divergent escape profiles (Figures 2C and 2D; Figure S2H). ØA antibodies (e.g., RSD5)^31^ target the β2–α1 loop and α4 helix, exhibiting exceptional potency (IC_50_ 5–16 ng mL^-1^) with slight RSV-B preference. ØB antibodies (e.g., CR9501, 5B11)^32,33^ depend more on the β2–α1 and β6–β7 loops, showing a 10-30-fold reduced neutralization without subtype bias with IC_50_ values of 127–137 ng mL^-1^ (Figures 2C–2E). ØC antibodies (e.g., MEDI8897) bind the α4 helix and achieve the strongest neutralization against RSV-A (IC_50_ ∼6 ng mL^-1^), but are stratified by sensitivity to RSV-B polymorphisms: the ØC_206/211_ subgroup (e.g., CNR2053; IC_50_ 484 ng mL^-1^) is impaired by I206M/S211N substitutions, whereas the ØC_205/208/209_ subgroup (e.g., MEDI8897; IC_50_ 452 ng mL^-1^) is sensitive to mutations at positions 208 and 209 (Figure S2I). Clinically, MEDI8897 exhibits a 10–1000-fold reduction against RSV-B strains carrying N208S/D^34^ or K209Q, and CNR2053 shows a ∼100-fold decrease against isolates with I206M/S211N^20,21,24^. In contrast, ØD antibodies (e.g., CNR3017) target cryptic residues N216 and E218, which are occluded in the prefusion state, rendering them non-neutralizing despite high affinity—a mechanism validated by cryo-EM (Figures 2C–2E; Figure S2J). Thus, site Ø identity alone does not predict antibody function, as neutralization potency varies by more than 1,000-fold across ØA to ØD.

Site V, centered on residue T174, partitions into two subclasses: VA (e.g., hRSV90)^35^, which engages hydrophobic residues A170, L172, S173, and A177 and extends toward site Ø, and VB (e.g., 01.4B)^36^, which primarily contacts V178 and D263 and orients toward site II (Figure 2C; Figure S2K). Both subclasses exhibit comparable, moderate neutralization (IC_50_ ∼75–185 ng mL^-1^), approximately 5–10-fold weaker than sites ØA/C antibodies. A very rare subset of VA antibodies (CNR2218, n=1 in pediatric-derived repertoires) cross-reacts with hMPV, mirroring the recently characterized RM 5-1 antibody, hinting at conserved structural motifs across hMPV^37^. Site III antibodies recognize a structurally conserved epitope spanning the β2 strand (T50–G51), α6–α7 helices (N262–K272), and β7 strand (G307–T311), which is conserved between RSV and hMPV (Figure 2B; Figures S2L–S2N). Consistent with this, approximately 47% of site III antibodies cross-neutralize hMPV (Figure S2O), a property shared with previously described broad-spectrum antibodies such as MPE8, RSV-199 and CNR2047^24,38,39^. Functional clustering based on DMS and neutralization profiles further divides site III into two subgroups: the III_hMPV-reactive_ subgroup engages the pan-pneumovirus-conserved β2/β7 patch (T50, G307, V308, D310, T311), tolerates variation at α7 (e.g., D269), and achieves increased potency via SHM-expanded interfacial contacts; in contrast, the III_RSV-specific_ subgroup binds closer to the genus-specific α7 segment (residues 266–269) and is sensitive to mutations such as D269 (Figures S2O and S2P). Site VI (e.g., 2.4K)^5,27^ is a membrane-proximal epitope; its key residues (D479, F477, I499) undergo substantial conformational changes during the preF-to-postF transition, rendering it structurally preF-specific (Figures 2A and 2B; Figure S2Q). However, most of the 22 site VI antibodies we identified (≈3% of the total repertoire) are non-neutralizing (Figure S2R). Structural analysis of two site VI antibodies, CNR2185 (this study) and 2.4K^5^, confirmed that their Fab constant domains orient downward toward the viral membrane, creating a steric conflict that likely limits neutralization potency (Figure S2S). Collectively, preF-specific binding is necessary but not sufficient for neutralization, with ØA/ØC as elite, ØB/V/III as moderate, and ØD/VI as non-neutralizing despite high preF affinity, defining an apical-to-basal potency gradient on preF.

### Cross-reactive and postF-specific epitopes define non-productive landscapes

Parallel analysis of cross-reactive and exclusively postF-specific epitopes identifies distinct antigenic regions that predominantly elicit non- or weakly neutralizing responses, providing a critical negative correlate for vaccine design to mask or disfavor these sites and focus immunity toward protective antibodies. Site II antibodies target a well-defined epitope^40,41^ centered on residue N268, with key contacts at N262, D269, K272, and S275 (Figure 3A; Figures S3A–S3C). Clinically observed mutations (e.g., N262D, K272E, S275F/L) and engineered variants (e.g., N268I, K272N/T/M/Q) at these positions abolish or reduce binding to therapeutic antibodies such as palivizumab^42-44^ confirming the functional relevance of this interface. The 40 site II antibodies identified in this study exhibit moderate neutralization potency, with geometric mean IC_50_ values of 268 ng mL^-1^ against RSV A2 and 449 ng mL^-1^ against B9320 (Figure 2A; Figure S3D). Site IV displays pronounced functional heterogeneity, dividing into two DMS-defined subclasses (Figures 3A and 3B). The IVA subclass, centered on G446 and exemplified by the clinical antibody MK1654 (Clesrovimab)^9,11^, achieves high neutralization potency (IC_50_ < 100 ng mL^-1^) (Figure 3C). Notably, the ten most potent IVA antibodies (IC_50_ < 10 ng mL^-1^) are cross-reactive with both preF and postF (Figure 3C), challenging the paradigm that only preF-specific locking confers elite neutralization. The IVB subclass, centered on G430, shows weaker neutralization (IC_50_ ∼600 ng mL^-1^) and further splits into RSV-only and hMPV-only neutralizers: RSV-specific IVB antibodies rely on R429 (mutated to V429 in hMPV), while hMPV-specific IVB antibodies target the conserved I431 (Figures 3D–3F). Non-neutralizing IVB antibodies engage an expanded footprint around G430 (K421, T434, S436), representing a distinct, non-functional binding mode (Figures 3D–3F). Together, these illustrate how subtle differences in residue preference and epitope topology within a single antigenic site dictate neutralization breadth, potency, and escape potential. Site I antibodies engage a structurally rigid and conformationally conserved epitope (N380–K390) spanning the pre-and post-fusion states (Figure 3A; Figure S3E). Cryo-EM analysis of CNR2358 bound to postF confirms that residues P389 and K390 constitute critical binding determinants (Figure S3F). Although most site I antibodies are cross-reactive, infant-derived ADI-14359 exhibits strict postF selectivity mediated by a single hydrogen bond to K390, illustrating that site I also contains discrete residues capable of driving conformation-specific recognition^45^. The preferential binding of site I antibodies to postF correlates with their limited neutralization (Figure S3G), suggesting that epitope accessibility or structural dynamics in the post-fusion state constrains functional efficacy.

**Figure 3.**
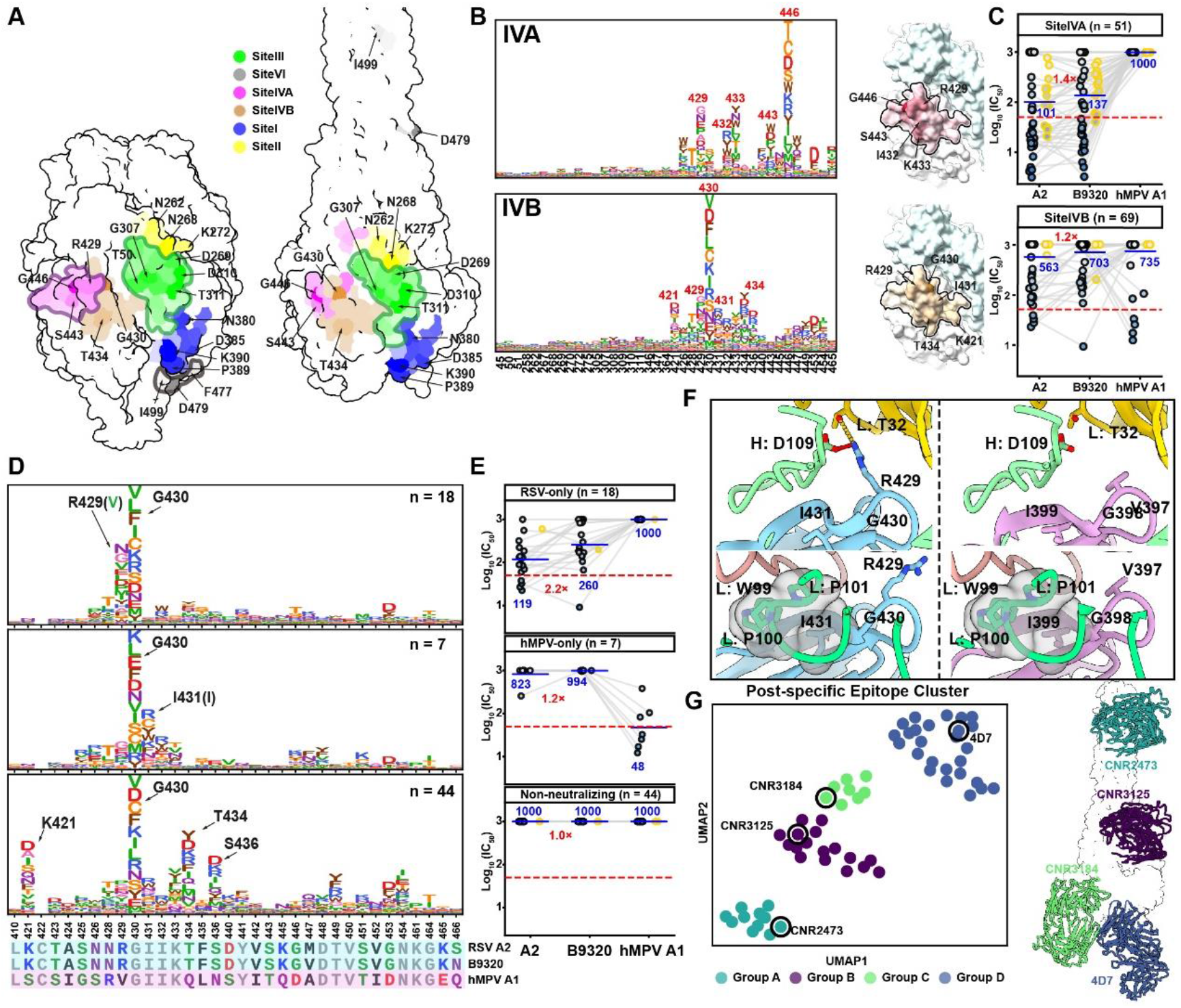
Structural and functional refinements of non-productive antibodies. **(A)** Surface representations of escape risk at sites II, III, IV, I, and VI on RSV preF and postF, colored as in Figure 2A. Arrows indicate critical escape residues, and color intensity reflects escape risk (darker, higher). PostF-specific antibody epitopes on RSV postF are also shown. **(B)** DMS escape profiles and structural projection of RSV F binding activity for site IVA and site IVB antibodies, with key residues labeled. **(C)** Neutralization potency (IC_50_) of site IV–targeting antibodies against RSV A2, B9320, and hMPV A1. **(D)** Escape mutation profiles of site IVB antibodies, including RSV-only neutralizers (n = 18), hMPV-only neutralizers (n = 7), and non-neutralizing antibodies (n = 44). Epitope amino acid multiple sequences alignment analysis for sites IV (lower panel). Conserved residues are shown in gray. Variable residues are colored by physicochemical properties: hydrophobic residues (A, L, I, P, F, and M), black; polar residues (G, S, and T), yellow; neutral residues (Q and N), purple; basic residues (K, H and R), blue; and acidic residues (D and E), red. **(E)** Neutralization potency (IC_50_) of site IVB–targeting antibodies against RSV A2, B9320, and hMPV A1. In (**C)** and (**E)** dot borders are colored by RSV F binding preference (pre-specific, yellow; cross-reactive, black). Blue lines and red values indicate geometric mean IC_50_ values of antibodies. Red dashed lines mark the threshold for very high neutralizing potency (50 ng mL^-1^ for site IV). **(F)** Structural analysis of representative antibodies CNR2298 (PDB ID: 27EN) and CNR2175 (PDB ID: 24JN) illustrates the structural basis for RSV-only or hMPV-only specificity. **(G)** Left: UMAP projection of RSV F postF–specific antibodies colored by epitope clusters. Each point represents a monoclonal antibody (mAb), with colors indicating distinct epitope groups defined by competitive binding assays combined with structural validation (four antigen–Fab complex structures determined in this study). Right: Binding positions of the four postF–specific epitopes.

We classified 62 postF-specific antibodies into four major groups (A–D) via BLI-based competition and provided their first structural characterization (Figure 3G). Group A (e.g., CNR2473) targets the exposed six-helix bundle via electrostatic and hydrogen-bond networks with α5 and α10 helices (Figure S3H). Group B (e.g., CNR3125) engages the β6–β7 loop through charged and polar interactions with residues including Ser290, Lys293, Glu294, and Glu295 (Figure S3I). Group C (e.g., CNR3184) binds near β22, forming extensive hydrogen-bond and salt-bridge contacts with residues from an adjacent protomer (Figure S3J). In contrast, Group D (e.g., 4D7)^46^ employs a distinct mode dominated by hydrophobic contacts with a patch formed by Leu381, Val384, Ile386, Phe387, and Leu334 (Figure S3K). Functionally, all postF-specific antibodies lack detectable neutralization. Mapping these non-productive sites provides a negative correlate for vaccine design that a high proportion of postF-specific or weakly neutralizing cross-reactive antibodies (sites I, II, IVB) signals misdirected immunity. In vaccine contexts, such responses may compete with protective apical targeting and, as discussed below, merit consideration in the context of VAERD.

### Divergent immunodominance hierarchies between infants and adults

Quantification of repertoire composition revealed starkly opposing immunodominance profiles (Figure 4A). In pediatricians, preF-specific antibodies constituted ∼75% of the repertoire, with high-potency clones targeting sites ØA–ØC, VA–VB, and IVA alone accounting for ∼55% (Figure 4A). In toddlers, although preF-specific antibodies still represented ∼55% of the repertoire, high-potency binders against these same key sites comprised only ∼20%, and critically, this 20% was predominantly composed of second-tier antibodies: the ØB and IVA classes, which exhibit compromised neutralization with IC_50_ > 200 ng mL^-1^ (Figures 4A and 4B). Most strikingly, top-tier ØA/ØC-specific antibodies constituted ∼23% of the pediatrician repertoire but were nearly undetectable in toddlers (<1%) (Figure 4B). Spatially, ∼80% of toddler-derived antibodies targeted the central-to-basal region (sites III, IV, I, and VI), most displaying marginal potency (Figures 4A and 4B). In contrast, pediatricians with repeated exposure showed a distinct bias toward upper, central-head sites, dominated by site Ø (38%), site V (17%), and site III (17%)—all associated with high neutralization activity (Figures 4A and 4B). This spatial and qualitative disparity points to fundamentally different epitope-activation thresholds between primary and mature immune responses.

**Figure 4.**
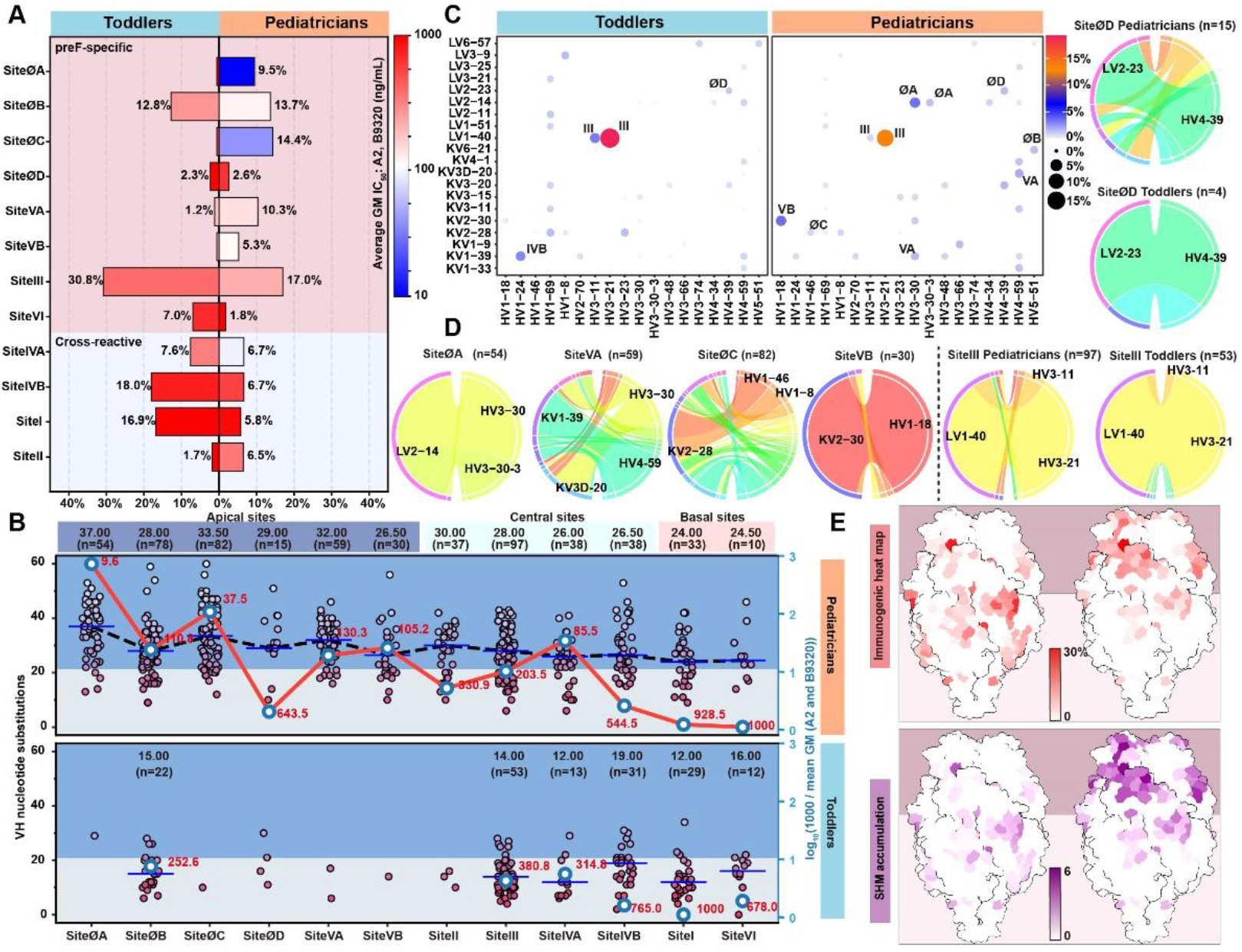
Distinct epitope targeting and germline usage in pediatrician and toddler RSV F antibody repertoires. **(A)** Proportion of antibodies across DMS-defined epitope groups in pediatrician and toddler cohorts. Mirror bar plots show the percentage of antibodies in pediatricians (right) and toddlers (left). Bar colors represent the log_10_ geometric mean IC_50_ against RSV A2 and B9320, with warmer colors denoting higher IC_50_ values. **(B)** Relationship between SHM and neutralization potency across DMS-defined epitope groups. Each point represents a monoclonal antibody colored by the number of VH nucleotide substitutions. Dashed lines indicate median values within each group, with sample sizes shown above. The red values represent neutralization potency calculated as log_10_ (1000 / mean GM) of RSV A2 and B9320. Higher vertical position of the blue dots indicates stronger neutralization. **(C)** Dot plot of VH/VL gene usage in RSV F-specific memory B cells from pediatricians and toddlers. VH/VL gene pairs used by < 1% of antibodies in all groups were omitted. **(D)** Chord plots showing VH/VL germline pairing of antibodies targeting sites ØA, ØC, VA and VB in pediatricians and sites ØD and III in both groups. Each chord represents the frequency of VH/VL germline combinations. Germline genes are labeled. **(E)** Surface representations of DMS-defined epitope intensity and VH SHM accumulation in toddler (left) and pediatrician (right) cohorts. Values were projected onto the RSV preF structure as described in Methods.

SHM analysis exposed a head-to-bottom decreasing gradient in adults (Ø, 32; V, 29.3; II, 30; III, 28; IV, 26; I, 24; VI, 24.5) (Figure 4B). Site ØB unusually showed lower SHM than neighboring apical sites, indicating a "low-threshold" window within an otherwise high-barrier region (Figure 4B). Sites III, IV, I, VI (collectively ∼85% of the toddler repertoire) and ØB (∼12%) are characterized by lower SHM requirements, indicating they are antigenically “permissive” and can be engaged rapidly by germline-encoded antibodies during primary infection (Figures 4B and 4C). Germline profiling revealed IGHV3-21/IGLV1-40 dominating site III across ages (linked to hMPV cross-reactivity), whereas adult-enriched high-potency specificities (ØA: IGHV3-30/IGLV2-14; VB: IGHV1-18/IGKV2-30) were virtually absent in toddlers (Figure 4C). VA and ØC antibodies, which are prevalent in adults but rare in toddlers, are preferentially encoded by IGHV4-59 and IGKV2-28, respectively (Figures 4C and 4D). Sites ØB, II, IVA, IVB, I and VI showed more diverse germline usage, suggesting weaker developmental constraints compared with site III and ØA (Figure S4A). Broader repertoire analysis showed that toddler repertoires were highly restricted, dominated by the IGHV3-21/3-11/IGLV1-40 pair (∼18.7% of all VH/VL combinations), whereas pediatrician repertoires displayed extensive germline diversity, enabling a broader array of high-potency antibodies (Figures 4C and 4D).

These patterns delineate two response modes: (i) an early, germline-encoded mode (sites III, IV, I) engaging low-SHM, permissive epitopes for rapid but suboptimal containment; (ii) a high-affinity maturation mode (ØA, ØC, V) requiring repeated exposure and extensive SHM. Pre-existing maternal antibodies against apical epitopes may further suppress infant priming to these sites, passively redirecting B-cell immunity toward subdominant, less protective specificities^47,48^. Repeated RSV exposure in pediatricians drives a marked shift in antibody distribution and quality, characterized by epitope redirection from basal to apical regions, progressive SHM accumulation, and enhanced neutralization potency (Figures 4B and 4E). A parallel dynamic occurs in influenza, where haemagglutinin site Cb-reactive antibodies dominate initial responses using a single gene pairing with minimal SHM, before being superseded by site Sb-reactive antibodies upon affinity maturation^47-49^. The immunodominance of readily accessible, low-SHM epitopes in primary infant responses provides a plausible mechanistic link to the failures of pediatric RSV vaccines, including VAERD. Whether site III/I/VI antibodies directly mediate enhanced pathology (via Fc or otherwise) remains to be tested in dedicated animal models, our data provide the immunological substrate for such tests but do not directly demonstrate pathology enhancement.

### Distinct maturation paths encode the threshold divide: site III vs. ØA

To dissect molecular basis of divergent thresholds, we traced two dominant lineages: IGHV3-21/IGLV1-40 (site III) and IGHV3-30/IGLV2-14 (ØA). Analysis of public antibody databases^50^ (>20,000 antibodies across nine viruses) revealed distinct germline biases: the IGHV3-21/IGLV1-40 pair was enriched specifically in RSV- and hMPV-targeting antibodies; whereas IGHV3-30/IGLV2-14 was broadly distributed across multiple viruses (Figure 5A).

**Figure 5.**
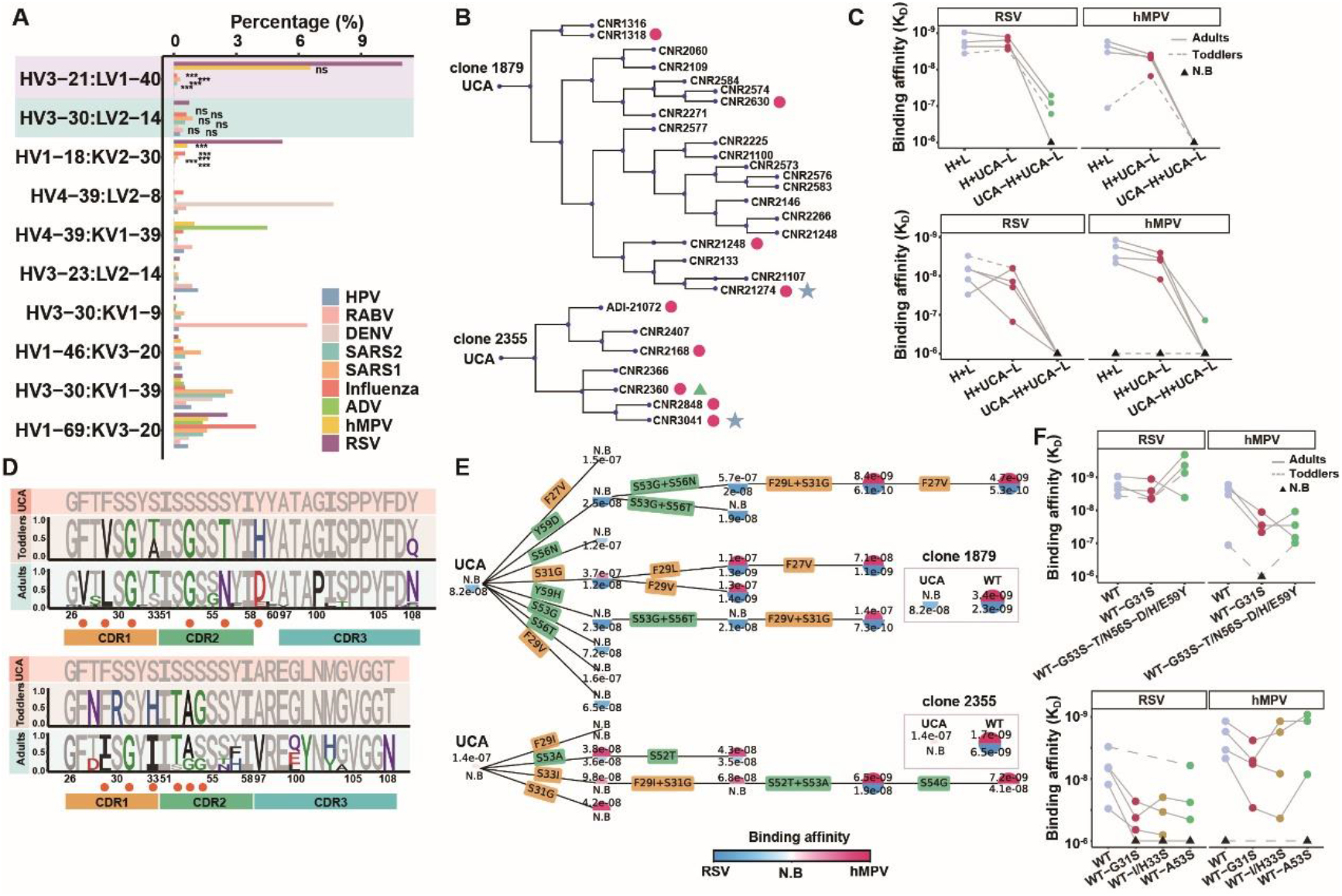
Affinity maturation pathways of IGHV3-21/IGLV1-40 encoded site III antibodies. **(A)** Bar plot showing the proportions of representative VH/VL pairs among nine viruses. Statistical significance compared with RSV was determined using Fisher’s exact test. ***p < 0.001; ****p < 0.0001; ns, not significant. **(B)** Genealogical trees of the IGHV3-21/IGLV1-40 antibody linage. Red dots indicate representative antibodies selected from each clone based on HCDR3 degeneracy for functional testing in Figure 5C. The blue stars denote antibodies subjected to additional substitution analysis in Figure 5C. The green triangle denotes the antibody whose light chain was used as a common pairing partner for clones 1879 and 2355 in Figure 5C. **(C)** Binding affinities of three antibody configurations against RSV and hMPV for clone 1879 (top) and clone 2355 (bottom): the fully mature antibody (H+L), the mature heavy chain paired with the unmutated common ancestor light chain (H+UCA-L), and the fully germline antibody (UCA-H+UCA-L). N.B., non-binding. **(D)** Sequence logo plots of CDR1/2/3 from clone 1879 (top) and clone 2355 (bottom), including the flanking residues adjacent to CDR2. Red dots indicate sites selected for mutagenesis screening. **(E)** Binding affinities of mutagenesis screening variants for clone 1879 (top) and clone 2355 (bottom). **(F)** Binding affinities of reversion mutants derived from clone 1879 (top) and clone 2355 (bottom). WT: the fully mature antibody. N.B., non-binding.

We reconstructed genealogical trees from combined datasets for two representative IGHV3-21/IGLV1-40 clones (1879 and 2355). These clones contained antibodies derived from both adults and infants/toddlers, and exhibited cross-reactivity against hMPV (Figure 5B). Focusing on heavy-chain contributions, we generated antibody combinations representing mature, partially reverted, and fully germline states. Binding assays showed that reverting the light chain to its unmutated common ancestor (UCA) caused only minor reductions (1- to 4-fold), whereas reverting the heavy chain to UCA led to a drastic 50–1000-fold loss of binding to both RSV and hMPV, confirming that antigen recognition is primarily driven by heavy-chain maturation (Figure 5C). To identify the specific residues that drive binding, we performed systematic mutagenesis scanning across HCDR1 and HCDR2. For clone 1879, four HCDR1 (F27V, F29L/V, S31G) and five HCDR2 (S53G, S56T/N, Y59D/H) substitutions were individually introduced into the UCA version of antibody CNR21274, which exhibits weak RSV binding and undetectable hMPV reactivity (Figures 5D and 5E). The single substitution S31G was sufficient to confer hMPV reactivity, while Y59D/H or S31G enhanced RSV binding 4-8-fold (Figures 5D and 5E). Although other single mutations had limited effects, combinations in HCDR1 (e.g., F27V/F29L/S31G) further improved affinity, reaching levels only 2-5-fold lower than the fully matured control (Figures 5D and 5E). Multiple HCDR2 substitutions (e.g., S53G, S56T, Y59D/H) further increased RSV binding but did not improve hMPV affinity, underscoring the essential role of S31G in cross-genus recognition (Figure 5E). Combining HCDR1 and HCDR2 mutations achieved affinities comparable to the mature antibody, suggesting that framework somatic hypermutation contributes minimally to affinity maturation in this context (Figure 5E). Loss-of-function reversion confirmed that G31S significantly impaired binding, especially to hMPV (Figure 5F). In clone 2355, S31G and S33I enhanced hMPV binding, and S53A improved binding to both viruses (Figures 5D and 5E). Notably, S31G enhanced hMPV binding in both clones (Figures 5E and 5F), highlighting its conserved role in cross-reactivity. Site III thus achieves functional maturation via minimal key mutations (notably S31G), representing a low evolutionary barrier.

For the IGHV3-30/IGLV2-14 lineage, genealogical reconstruction revealed five distinct clonal families (Figure S5A). Five combinations per clone were generated for functional testing: (i) mature heavy chain (H) + unmutated common ancestor light chain (UCA-L), (ii) UCA-H + mature L, (iii) UCA-H + UCA-L, (iv) H with UCA-reverted HCDR1 + mature L, and (v) H with UCA-reverted HCDR2 + mature L (Figure S5B). We observed that all antibody combinations bearing at least one chain reverted to the UCA state lost RSV binding activity. Moreover, most antibodies lost binding when only HCDR1 was reverted, and nearly all lost binding upon HCDR2 reversion (Figure S5B). These indicate that high-affinity engagement of the ØA epitope requires a sophisticated, cooperative set of mutations across both chains and multiple CDR regions, reflecting a substantially higher evolutionary barrier.

In primary infection, infant responses are dominated by low-SHM, IGHV3-21-encoded antibodies targeting site III; only ∼2% (2/92) in infants^45^, 7% (3/44) in toddlers carry the S31G mutation, correlating with limited neutralization and minimal hMPV cross-reactivity. In contrast, the adult repertoire is enriched with high-SHM site III antibodies, ∼25% (6/25) in healthy adults^27^ and 34% (32/94) in pediatricians with S31G, enabling broad hMPV cross-neutralization, alongside high-potency ØA-specific antibodies bearing the highest SHM burden. These mechanistically explain why infants preferentially elicit abundant, marginally potent site III antibodies, while the ØA imposes a high evolutionary barrier that may not be surmounted without repeated antigen exposure, revealing this epitope-intrinsic maturation threshold as the primary driver of the well-documented infant-adult antibody repertoire gap.

### Adult-driven immune pressure shapes RSV antigenic evolution

To understand how these human antibody repertoires sculpt RSV evolution in nature, we integrated global genomic surveillance with functional antigenic mapping. We analyzed 27,013 RSV A and 21,174 RSV B high-quality genomes from GISAID, supplemented by 1,687 newly sequenced genomes (941 A, 746 B) from patients with acute respiratory tract infections (2009–2025) in China. Antigenic phylogenetic analysis of RSV F delineated six RSV A and six RSV B lineages with distinct antigenic signatures (Figures 6A and 6B; Figures S6A and S6B). Although antigenic drift in RSV A remains weak, with only two high-frequency substitutions at non-epitope sites (S276N and S377N), two novel RSV B lineages (clade 1 and clade 3) emerged around 2020 and rapidly achieved global dominance, now accounting for >90% of circulating strains (Figure 6A; Figure S6B). RSV B circulating lineages have progressively accumulated signature mutations within high-potency functional epitopes, primarily ØC and V (Figures 6B and 6C). Although substitutions such as S389P (site I) and K42R/S190N (outside known epitopes) are not themselves neutralizing targets, their tight linkage with ØC- and VA-associated mutations forms an epistatic cluster (Figure 6B). Key residues—L172, S173 and I206—exhibited progressive, lineage-replacing trajectories that have systematically displaced ancestral variants since 2009. In contrast, K209 showed a rare oscillating pattern with seasonal co-circulation of multiple variants, suggesting complex fitness trade-offs between immune escape and viral function (Figure 6C). Critically, since 2024 the S211N substitution has become the dominant variant (lineage RSVB25), frequently co-occurring with L172Q, S173L, I206M, K209R and the epistatically linked R42/S389 mutations, collectively amplifying immune evasion (Figures 6B and 6D). Functional assays using clinical and laboratory strains confirmed the escape potential of these mutations. ØC-specific antibodies suffered 20-fold and 25-fold neutralization losses against strains carrying K209Q (lineage B18537) and the I206M/K209R/S211N combination (lineage RSVB25), respectively (Figure 6D). In contrast, ØA antibodies retained broad-spectrum potency against all tested RSV-B strains. VA antibodies showed a 2–3-fold reduction from L172Q (RSVB63/71 isolates) and near-complete loss with combined L172Q/S173L mutations (Figure 6D; Figure S6C). These impairments align precisely with DMS-predicted escape profiles, validating the functional relevance of naturally occurring polymorphisms.

**Figure 6.**
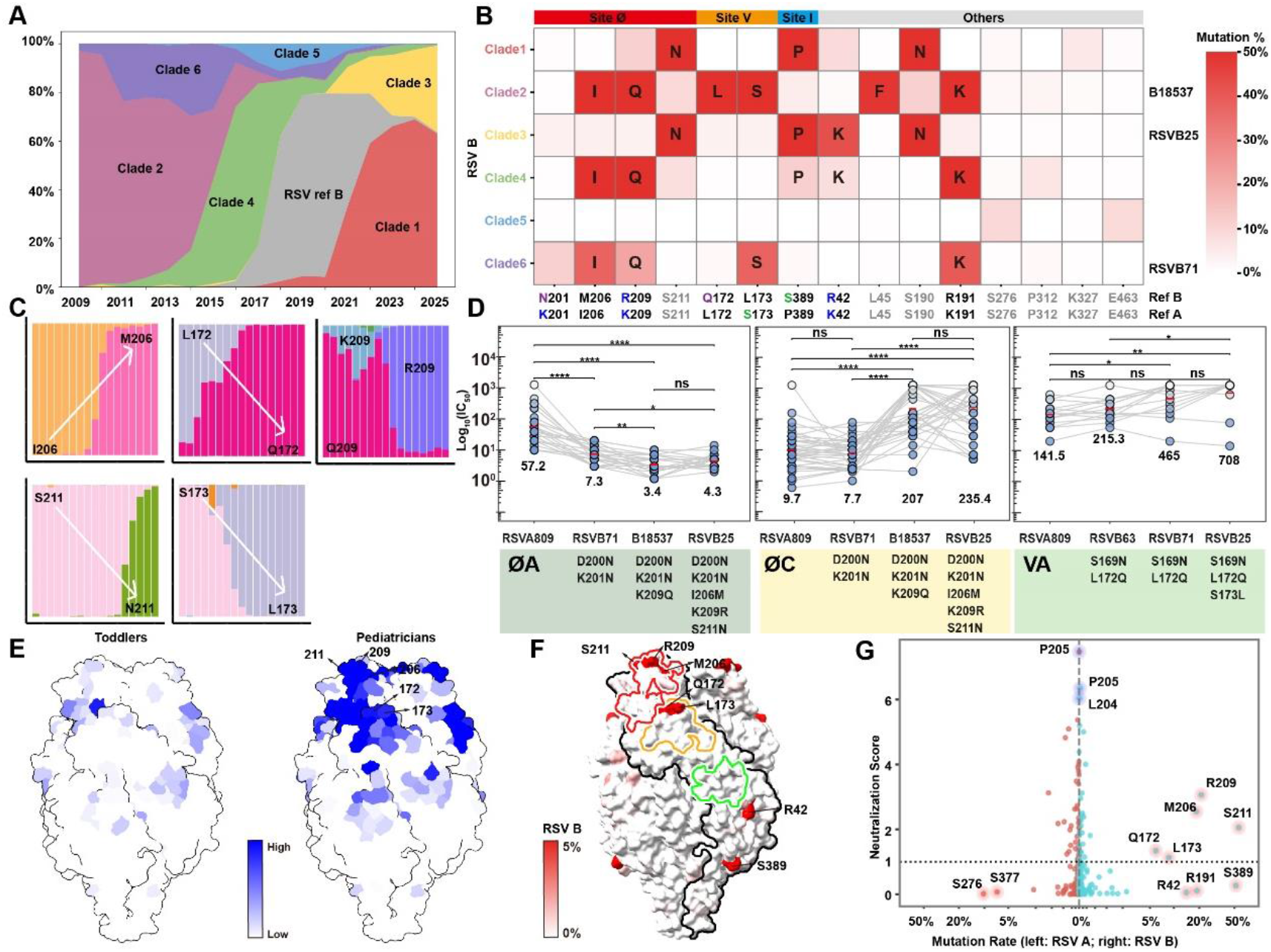
Immune pressure-driven antigenic evolution of RSV F. **(A)** Temporal dynamics of RSV-B clades and mutation frequency across key sites. Stacked area plot showing the yearly distribution of RSV-B clades from 2009 to 2025. Each color represents a distinct clade, while grey denotes wild-type (WT) strains. **(B)** Mutation rates of antigenic-site residues with overall mutation frequencies≥1% in RSV-B. Each residue is colored according to its mutation rate within the corresponding clade. **(C)** Temporal dynamics of representative single mutation sites in RSV-B. **(D)** Neutralization potency (IC_50_) of ØA-, ØC-, and VA-targeting antibodies against RSV clinical isolates containing critical escape mutations (as labeled below), with RSVA809 (subtype A) served as a control. Red dots and black values indicate the geometric mean IC_50_ of antibodies. **(E)** Surface representations comparing neutralization potency between toddler and pediatrician cohorts. Site-level neutralization potency was projected onto corresponding epitopes to quantify neutralization potency at each antigenic site. **(F)** Surface representations of RSV F based on publicly available sequences of RSV B. The color bar indicates mutation rate from 0% to 5%, with increasing red shading representing greater variability. Major antigenic regions are outlined in red for site Ø, orange for site V and green for site III. **(G)** Scatter plot showing the relationship between mutation rate (x-axis; RSV A on the left and RSV B on the right) and neutralizing potency. ***p<0.001, ****p<0.0001, ns, no significance.

Projecting functional epitope profiles from toddler- and adult-derived antibodies onto the preF structure, weighted by neutralization potency, allowed us to map the “neutralization intensity” exerted at each residue. Integrating global genomics (2009–2025) with functional mapping revealed that adult immune pressure focuses on apical sites ØA–ØC/V, while toddler pressure is weaker and centers on ØB/III (Figure 6E). Strikingly, RSV-B mutation hotspots (172, 173, 206, 209, 211) cluster precisely within ØC and V in which the epitopes under strongest adult neutralization pressure (Figure 6F). Residues 204–205, though under maximal predicted pressure, remain nearly invariant presumably due to structural constraints on α4–α5 (Figure 6G), creating a high-escape-barrier core that ØA antibodies exploit (ØA retained potency against all tested RSV-B strains)^24^. RSV-A showed weaker correlation between hotspots and neutralization intensity, suggesting subtype-specific fitness trade-offs. These indicate that repeatedly exposed adults, rather than infants, act as the dominant selective force shaping contemporary RSV (especially RSV-B) evolution, creating an adult-centric, dynamically shifting antigenic target that further disadvantages naive infants whose antibody repertoires are already restricted to low-threshold epitopes.

## Discussion

This study constructs a high-resolution, functionally annotated antigenic atlas of RSV F by integrating deep mutational scanning (DMS) with comparative repertoire profiling of repeatedly exposed pediatricians and RSV-experienced toddlers. Moving beyond canonical structural classification (sites Ø–VI), our data resolve 12 functional epitope subclasses, including four subcategories within immunodominant site Ø that span a 1000-fold potency gradient (Figure 2), and define a maturation-gated functional hierarchy of preF: apical epitopes (ØA, ØC, V) impose high activation barriers requiring extensive somatic hypermutation (SHM) and cooperative mutations across multiple complementarity-determining regions, whereas central-to-basal epitopes (III, IV, I) are permissively engaged by germline-encoded antibodies via minimal substitutions such as S31G (Figures 4 and 5). This threshold logic governs repertoire partitioning: toddler responses default to low-barrier, non-productive central-to-basal sites, whereas repeated occupational exposure in pediatricians drives a maturation-shifted immunodominance toward apical, high-potency targets, a skew likely reinforced by maternal apical antibodies that suppress infant priming to these sites^27,45,47,48^. Genomic surveillance further shows adult-derived neutralizing pressure on apical ØC/V constitutes the dominant driver of contemporary RSV-B evolution (Figure 6), with structural constraints at residues 204–205 creating a high-escape-barrier core that ØA antibodies exploit^24^. These data reframe pediatric RSV vulnerability as a threshold-gated repertoire deficit rather than a mere failure to engage preF.

The clinical dichotomy that stabilized preF vaccines protect adults yet fail or show insufficient efficacy in infants with some candidates linked to VAERD has long lacked an antibody-repertoire-level mechanistic explanation. Our study provides key mechanistic insights: infants primed on low-threshold, non-productive epitopes (III, I, VI, ØB) generate qualitatively suboptimal responses regardless of preF antigen delivery, as the maturation barrier to apical, protective epitopes (ØA/ØC/V) cannot be surmounted without repeated antigen exposure or tailored immunogen design. This reframes the pediatric RSV vaccine problem from "induce preF-binding antibodies" to "redirect immunodominance toward high-barrier apical targets while suppressing low-threshold permissive epitopes". Promising strategies may include (i) structural masking of subdominant, basally exposed non-neutralizing epitopes; (ii) epitope-focused immunogens that lower B-cell precursor activation thresholds for ØA/ØC; (iii) adjuvant and delivery platforms that promote robust early-life germinal center reactions and somatic hypermutation. Monitoring vaccine-induced repertoires for low frequencies of postF-specific and other non-productive antibodies may serve as a measurable negative correlate of protection.

Admittedly, our data do not directly test whether site III-, I-, or VI-targeting antibodies mediate VAERD. The current cohorts comprise naturally infected toddlers, rather than infants subjected to the formal VAERD paradigm of vaccine priming followed by RSV infections; direct evidence for pathology mediation (via FcγR engagement, complement activation, eosinophilic infiltration, etc.) will therefore require dedicated infant animal models. That said, the immunodominance of low-potency, germline-encoded antibodies at these central-to-basal, low-threshold epitopes in primary infant responses provides a concrete mechanistic substrate for VAERD interrogation: our functional atlas specifies which antibody specificities should be prioritized for passive transfer or vaccine-priming validation in relevant models, converting a previously descriptive "suboptimal infant humoral response" hypothesis into a testable, epitope-resolved framework.

This study has several limitations. First, while extensive, the repertoire analysis is derived from specific cohorts (Chinese pediatricians and toddlers); validation in broader geographic and ethnic populations is warranted. Second, the functional characterization of antibodies primarily relies on *in vitro* neutralization; their potential roles in Fc-mediated effector functions or immunopathology *in vivo* remain to be fully elucidated^51,52^. Third, while we delineate B cell-intrinsic mechanisms, the roles of T cell help, regulatory cells, and the mucosal environment in shaping these responses^53-56^ warrant further investigation. Fourth, the infant cohort provides a snapshot of natural infection; longitudinal studies tracking repertoire evolution from birth through repeated exposures are needed to fully chart the maturation trajectory. Finally, the proposed vaccine design principles require empirical testing in animal models and clinical trials. In summary, this study shifts the RSV vaccine paradigm from quantity (preF-binding titer) to quality (epitope specificity, potency, maturation state) governed by threshold. By mapping infant vulnerability and adult protection onto a functional epitope landscape with a maturity-gated hierarchy, we provide a framework, not yet a solution, for rationally designed pediatric RSV vaccines that bridge the immunity gap.

## Experimental model and subject details

### Human Subjects

Adult donors were healthy pediatricians working in the Department of Respiratory Medicine at the Children’s Hospital of Chongqing Medical University and had no respiratory infection symptoms within one month prior to enrollment. Infant and toddler donors were recruited from the same hospital without acute infection at the time of sampling. All toddlers had a documented history of natural RSV infection. Written informed consent was obtained from all adult participants and from the legal guardians of infant and toddler participants. Peripheral blood samples were collected by trained medical staff following standard procedures. The study was approved by the Ethics Committee of the Children’s Hospital of Chongqing Medical University, Chongqing, China.

## Method details

### Cells and viruses

HEK293F cells (Thermo Fisher Scientific) were cultured in OPM-293 CD05 medium (OPM Biosciences) in a shaker incubator at 37 °C with 8% CO₂. HEp-2 cells (ATCC CCL-23) and Vero E6 cells were maintained in Dulbecco’s modified Eagle medium (DMEM) supplemented with 10% fetal bovine serum (FBS). Respiratory syncytial virus (RSV) strains A2 (VR-1540) and B9320 (VR-955) were obtained from the American Type Culture Collection (ATCC). RSV clinical isolates were passaged on HEp-2 cells prior to use in neutralization assays. RSV were propagated in HEp-2 cells in DMEM supplemented with 2% FBS and purified by density gradient centrifugation. Human metapneumovirus (hMPV) was kindly provided by Dr. Yao Zhao (Children’s Hospital of Chongqing Medical University, Chongqing, China). hMPV was propagated in Vero E6 cells cultured in DMEM supplemented with 2% FBS and 0.00025% trypsin, and purified by density gradient centrifugation.

### Expression and Purification of RSV F and hMPV F Proteins

The prefusion-stabilized RSV F protein (DS-Cav1) containing residues 1–513 and four stabilizing mutations (S155C, S190F, V207L, and S290C) was cloned into the pCAGGS expression vector^16^. The construct included a C-terminal T4 fibritin trimerization motif and an 8×His tag to promote trimer formation and facilitate purification. A monomeric prefusion RSV F construct lacking the T4 fibritin trimerization motif was also generated. A postfusion RSV F construct containing the ectodomain of RSV F (ACO83301.1) followed by a 6×His tag was similarly prepared. The hMPV F (A1 subtype) ectodomain (residues 1–490) containing stabilizing mutations (A185P, A113C, and A339C)^57^ was also cloned into pCAGGS, along with a monomeric version lacking the trimerization motif. For protein production, plasmids encoding the respective constructs were transiently transfected into HEK293F cells cultured in suspension at 37 °C with 8% CO₂ and shaking at 130 rpm. Supernatants were harvested 72 h post-transfection by centrifugation at 1,000 × g for 30 min to remove cells and debris. Recombinant proteins were purified from clarified supernatants by Ni–NTA affinity chromatography, followed by size-exclusion chromatography on a Superdex 200 column (GE Healthcare) equilibrated in 20 mM Tris-HCl (pH 8.0) and 200 mM NaCl.

### PBMC isolation and Single B cell sorting

Plasma was removed by centrifugation at 2,000 rpm for 10 min. The cellular fraction was diluted and layered onto Ficoll-Paque PLUS (Cytiva) for density gradient centrifugation. The PBMC layer was collected, washed with calcium- and magnesium-free D-Hanks buffer, and treated with ACK lysis buffer if necessary to remove residual red blood cells. The mixture of RSV preF and postF protein was fluorescently labeled with Alexa Fluor 488 (BioLegend). PBMCs were stained using anti-human IgG (PE), IgD (PerCP/Cy5.5), and IgM (PerCP/Cy5.5), CD19 (PE/Cy7), CD27(APC). CD19⁺IgD⁻IgM⁻IgG⁺CD27⁺ B cells binding to RSV F protein were single-cell sorted into 96-well PCR plates using a FACSAria III flow cytometer (BD Biosciences).

### Single B cell cloning

Antibody variable genes (IgH, Igλ and Igκ) were amplified from single B cells by RT–PCR. For a subset of antibodies, paired heavy- and light-chain variable-region sequences were obtained from single-cell V(D)J sequencing using the 10x Genomics platform^58^. The amplified or sequence-derived variable regions were assembled with human IgG1 constant regions into transcriptionally active DNA (TAD) expression cassettes containing a CMV promoter and poly(A) signal using overlap extension PCR. TADs encoding paired heavy and light chains were co-transfected into HEK293T cells for transient antibody expression. Cell culture supernatants were harvested 72 h after transfection and screened for RSV binding. Positive clones were sequenced, and variable regions were cloned into pCAGGS expression vectors carrying human IgG1 constant regions for recombinant antibody production.

### Expression and purification of IgGs and Fab fragments

The variable region sequences of candidate antibodies were synthesized and cloned into pcDNA3.1 expression vectors containing the human IgG1 constant region. Recombinant heavy- and light-chain plasmids were co-transfected into HEK293F suspension cells at a 1:1 ratio and cultured at 37 °C with 8% CO₂ and shaking at 130 rpm for 5-7 days. Culture supernatants were collected by centrifugation at 4 °C to remove cell debris and filtered through a 0.45 μm membrane. Antibodies were purified by Protein A affinity chromatography. After washing with PBS, bound antibodies were eluted using acidic glycine buffer (0.1 M, pH 2.5-2.8) and immediately neutralized with Tris buffer (1 M, pH 8.0-9.0). Purified antibodies were concentrated and buffer-exchanged into phosphate-buffered saline (PBS, pH 7.2) using ultrafiltration units (100 kDa cutoff).

### Biolayer interferometry binding analysis

Binding affinities between antibodies and RSV preF or hMPV preF proteins were measured by biolayer interferometry (BLI) using an Octet system. Purified antibodies (20 μg mL^-1^) were immobilized on Protein A biosensors. Sensors were equilibrated in running buffer to establish a baseline, followed by antibody loading and a second baseline step. Antibody-loaded sensors were then transferred to wells containing RSV preF or hMPV preF (starting at 100 nM) to monitor association, and subsequently moved to buffer-only wells to record dissociation. A blank reference was included for background subtraction. Sensors were regenerated between cycles using glycine (pH 1.5) and re-equilibrated in running buffer. Binding curves were fitted using the Octet Data Analysis software and the data were fit to a 1:1 binding model to calculate association and dissociation rate constants, and *K*_D_ was calculated using the ratio *k*_d_/*k*_a_.

### BLI competition analysis, clustering, and visualization of postfusion-specific antibodies

Competition binding assays of postfusion-specific antibodies were performed using biolayer interferometry (BLI) on an Octet system (ForteBio). RSV postfusion F protein containing a C-terminal His tag was immobilized on Ni-NTA biosensors, followed by sequential association of a primary antibody and a secondary antibody. Competition responses were analyzed using ForteBio Data Analysis software to determine binding interference between antibody pairs.

Competition measurements for 62 monoclonal antibodies (mAbs) together with five reference antibodies (4D7, CNR2472, CNR2473, CNR2475, and CNR3184) were compiled into a competition matrix in which antibody pairs were represented as indices. Negative or missing values were replaced with zero. The matrix was then transposed so that each antibody was represented as a feature vector describing its competition profile across other antibodies. The data were standardized using z-score normalization prior to dimensionality reduction. Uniform Manifold Approximation and Projection (UMAP, n_neighbors = 8, min_dist = 0.8) was applied to project the competition profiles into two-dimensional space. Antibody clusters were subsequently identified using agglomerative hierarchical clustering with Ward linkage, yielding four clusters. The resulting embeddings were visualized using scatter plots with cluster assignments indicated by color. Antibodies with cryo-electron microscopy (cryo-EM) structures (4D7, CNR3184, CNR3125, and CNR2473) were specifically annotated to facilitate structural interpretation.

### Enzyme-linked immunosorbent assay (ELISA)

Purified RSV preF or hMPV preF proteins were diluted to 2 μg mL^-1^ in PBS and coated onto 96-well plates (100 μL per well) overnight at 4 °C. Plates were washed three times with PBST (PBS containing 0.05% Tween-20) and blocked with 1% BSA for at least 2h at room temperature. Antibodies were diluted to 1 μg mL^-1^ and serially diluted before being added to the plates (100 μL per well) and incubated for 1h at room temperature. After washing 3-5 times with PBST, HRP-conjugated goat anti-human IgG secondary antibody (Sigma-Aldrich) diluted 1:5,000 was added and incubated for 1 h at room temperature. Plates were washed again, followed by addition of TMB substrate (100 μL per well) for ∼10 min in the dark. The reaction was stopped with 2 M H₂SO₄, and absorbance was measured at 450 nm.

### Antigen–antibody complex structural analysis

Structural epitope and paratope contacts were defined from antibody–RSV F complex structures using a distance-based approach. For each PDB structure^5,10,11,16,24,27-33,35-37,39-41,58-66^, antigen and antibody heavy- and light-chain identifiers were manually assigned. Non-hydrogen atoms were parsed from ATOM records, and pairwise Euclidean distances were calculated between antigen atoms and antibody atoms. Antigen–antibody atom pairs within 4.5 Å were considered contacts. Atom-level contacts were collapsed to unique residue-level antigen–antibody contact pairs. Antigen residues contacting either the heavy or light chain were defined as structural epitope residues, and the corresponding antibody residues were recorded as paratope residues. Structural epitope residues were mapped to predefined RSV F antigenic regions: site Ø, residues 63–96 and 195–227; site V, residues 55–62, 146–194 and 287–300; site II, residues 254–277; site III, residues 46–54, 301–312, 345–352 and 367–378; site IV, residues 422–471; site I, residues 26–45, 313–319 and 379–390; and site VI, residues 473–513. Residues outside these regions were classified as undefined. For each antibody, the fractional epitope composition was calculated as the proportion of unique epitope residues falling within each antigenic region. To compare structural footprint size across antibodies, epitope coverage was further normalized by the length of the corresponding antigenic region. For antibodies with established site assignments, the assigned site was used for normalization; otherwise, the site containing the largest number of contacted residues was used as the dominant site.

### Construction of RSV F mutant libraries

To systematically evaluate antibody escape, we constructed two deep mutational scanning (DMS) libraries: a mammalian cell-expressed trimeric prefusion-stabilized RSV F (DS-Cav1) library and a yeast surface-displayed RSV F monomer library. For the mammalian system, mutagenesis was performed on the DS-Cav1 protein (residues Q26–G517), which incorporates four stabilizing mutations (S155C, S190F, V207L, S290C) to maintain the prefusion trimer conformation^16^. For yeast surface display DMS, an RSV F monomer construct was designed following previous work^24-26^. The construct contained residues Q26–A504, wherein the furin cleavage sites and a non-epitope region (R109–V144) were replaced by a flexible GGSGGS linker to maintain structural stability. Briefly, as detailed above, targeted NNS mutagenesis at key epitope positions and subsequent tagging with unique 26-nucleotide barcodes were performed via two rounds of PCR for both constructs. The resulting plasmid libraries were subjected to PacBio long-read sequencing to establish reliable barcode-mutation associations.

Stable mammalian cell lines expressing the DS-Cav1 mutational library were generated following a previously described protocol^67^. Barcode-linked mutant fragments were cloned into the lentiviral vector pLVX-T2A-ZsGreen. Lentiviral particles were produced in HEK293T cells by co-transfection with psPAX2 and pMD2.G packaging plasmids. Virus-containing supernatants were collected and used to transduce HEK293T cells at a low multiplicity of infection (MOI = 0.1) to ensure single viral integration events. Transduced cells were then selected with puromycin (10 μg mL^-1^) to generate stable cell populations expressing RSV F variants. For the yeast library, the resulting mutant fragments were cloned into the pETcon vector and introduced into Saccharomyces cerevisiae EBY100 cells via lithium acetate transformation. Following initial selection on SD-CAA agar plates, the transformants were expanded in liquid SD-CAA and subsequently induced for surface expression in SG-CAA medium.

### Antibodies escape cell sorting

Antibody escape profiling was performed using fluorescence-activated cell sorting (FACS). For mammalian libraries, HEK293T cells expressing RSV F variants were incubated with monoclonal antibodies at 4 °C for 30 min in FACS buffer. Cells were stained with Alexa Fluor 647-conjugated anti-c-Myc antibody to monitor surface expression and PE-conjugated anti-human IgG Fc antibody to detect antibody binding. Cells expressing RSV F (ZsGreen⁺Myc⁺) but exhibiting reduced antibody binding (PE⁻) were sorted as potential escape variants. Sorted cells were lysed by heating at 72 °C for 3 min, followed by immediate cooling on ice. Reverse transcription was performed in a 10 µL reaction containing HiScript II reverse transcriptase, RNase inhibitor, 5× buffer, DTT, betaine, MgCl_2_, and TSO. After incubation at 37 °C for 60 min, the reaction was inactivated at 85 °C for 5 s. The N26 barcode region was PCR-amplified from the resulting cDNA, purified using AMPure XP beads, and subjected to next-generation sequencing.

For the yeast display library, induced cells were incubated with primary antibodies followed by fluorescent secondary antibodies (anti-HA-APC, anti-Myc-FITC, and anti-human IgG-PE). Yeast cells positive for surface expression but showing reduced antibody binding were sorted by FACS. Recovered yeast populations were washed, regrown overnight, and plasmids were extracted. The N26 barcode region was PCR-amplified, purified using AMPure XP beads, and subjected to high-throughput single-end sequencing.

### DMS data analysis and antibody clustering

Following established DMS analysis frameworks^25,26^, escape scores were computed after processing sequencing reads and quantifying barcode counts in both the reference and antibody-selected populations via the dms_variants package (v1.6.0)^25^. Specifically, the calculation was defined as: Escape score=F × (n_X,ab_ /N_ab_)/ (n_X,ref_ /N_ref_). Here, n_X,ab_ and n_X,ref_ represent the specific counts of variant X within the antibody-selected and reference libraries, respectively; N_ab_ and N_ref_ denote the total barcode counts in each sample; and F is a normalization factor defined as the 99th percentile of escape ratios.

For antibody clustering and visualization, we used graph-based unsupervised embedding approaches. First, site escape scores, defined as the sum of mutation escape scores per residue, were normalized for each antibody so that the total sum across all residues was 1, representing a probability distribution over the RSV F surface. The pairwise dissimilarity between antibodies was computed using the square root of the Jensen–Shannon divergence (JSD) between their normalized site escape profiles, implemented via scipy.spatial.distance.jensenshannon (v1.7.0). A k-nearest-neighbor (k-NN) graph (k=15) was then constructed using python-igraph (v0.9.6), followed by Leiden clustering to assign each antibody to an epitope group^68^. Cluster identities were manually annotated based on shared escape features within each group. To visualize antibody relationships in two dimensions, we applied UMAP using the umap-learn module (v0.5.2). Final plots were generated in R using the ggplot2 package (v3.3.3).

### DMS-defined antigenic-site assignment and site-level functional annotation

DMS-defined antigenic-site assignments were derived from antibody-specific escape profiles. Mutation escape scores were first summed at the antigenic-site level to generate an antibody-by-site escape matrix. For each antibody, the maximum site-level escape score was identified, and sites with scores greater than or equal to 10% of this maximum were retained as DMS-supported sites. Retained sites were ranked in descending order of escape magnitude, and up to the top ten sites were assigned to each antibody. This approach permitted antibodies with broad or composite escape profiles to be assigned to multiple antigenic sites, while minimizing contributions from low-level background escape signals. The same assignment strategy was applied to preF-specific, cross-reactive, pediatrician-derived and toddler-derived antibody subsets.

Site-level neutralization potency was estimated by aggregating reciprocal IC_50_ values across antibodies assigned to each DMS-defined site. For each antibody, IC_50_ values against RSV A2 and RSV B9320 were converted to reciprocal potency scores, with non-finite, missing or non-positive values assigned a score of zero. For each antigenic site, reciprocal IC_50_ values were summed across all assigned antibodies separately for RSV A2 and RSV B9320. The mean site-level neutralization potency was defined as the average of the two strain-specific reciprocal IC_50_ scores. Antibodies assigned to multiple sites contributed once to each assigned site. To enable comparison across cohorts with different numbers of antibodies, site-level neutralization potency values were normalized by the total number of antibodies in the corresponding cohort or antibody population.

Site-level VH somatic hypermutation accumulation was calculated using VH nucleotide substitution counts. For each antibody, VH mutation burden was defined as the sum of heavy-chain nucleotide mismatches and gaps. Missing or non-finite values were set to zero. For each DMS-defined antigenic site, VH mutation counts were summed across all antibodies assigned to that site. Antibodies assigned to multiple DMS-defined sites contributed their VH mutation count once to each assigned site. To enable comparison between antibody populations with different sample sizes, site-level VH SHM accumulation values were normalized by the total number of antibodies in the corresponding population.

### RSV Indirect immunofluorescence assay

Neutralization activity of monoclonal antibodies (mAbs) against RSV laboratory strains A2 and B9320 was determined by an indirect immunofluorescence assay. HEp-2 cells were seeded in 96-well flat-bottom plates and incubated overnight at 37 °C with 5% CO_2_. Antibodies were serially diluted threefold (8 dilutions) in culture medium. An equal volume of RSV containing 200 plaque-forming units (PFU) per well was mixed with diluted antibodies and incubated for 1 h at 37 °C with 5% CO_2_. The antibody–virus mixtures (100 μl per well) were then added to HEp-2 cells and incubated for 1 h at 37 °C to allow viral adsorption. Following infection, cells were overlaid with DMEM supplemented with 2% FBS and 1.2% methylcellulose and incubated for 22 h at 37 °C with 5% CO_2_. Cells were subsequently fixed with 80% acetone in PBS at −20 °C for 20 min and blocked with 2% bovine serum albumin (BSA) for 30 min at room temperature. RSV-infected cells were detected using a primary antibody followed by Alexa Fluor 488–conjugated goat anti-human IgG secondary antibody. Plates were washed three times with PBST (PBS containing 0.05% Tween-20) prior to secondary antibody incubation and five times after incubation. Fluorescent signals were measured using an automated spot reader (CTL Analyzer LLC) in the fluorescein isothiocyanate (FITC) channel. Neutralization curves were generated by nonlinear regression, and the half-maximal inhibitory concentration (IC_50_) values were calculated using GraphPad Prism (v10.1.2). An inter-assay normalization factor was calculated by averaging the IC_50_ of reference control (MEDI8897) across all batches. Normalized IC_50_ values were then calculated as (raw IC_50_ / batch-specific MEDI8897) × mean MEDI8897 across batches. Final normalized values above 1000 ng mL^-1^ were capped at 1000 ng mL^-1^.

### Fluorescence-based hMPV neutralization assay

Neutralizing activity against human metapneumovirus (hMPV) was measured using a fluorescence-based neutralization assay. Antibodies were serially diluted in hMPV maintenance medium consisting of DMEM supplemented with 3% FBS and 0.0025% trypsin. Diluted antibodies, negative controls and positive reference antibodies were prepared in 96-well plates, with duplicate wells for each dilution. Cell-control and virus-control wells were included on each plate.

An equal volume of diluted hMPV was added to antibody-containing wells and virus-control wells, whereas maintenance medium was added to cell-control wells. Antibody–virus mixtures were incubated at 37 °C with 5% CO_2_ for 1–2 h. Vero E6 cells in good growth condition were then trypsinized, resuspended in hMPV maintenance medium and added to each well at 3–4 × 10^5^ cells mL^-1^, 100 μL per well. Plates were incubated at 37 °C with 5% CO_2_ for 24 ± 2 h. After incubation, plates were imaged using a high-throughput fluorescence microplate imaging system, and fluorescent foci were counted for each well. The percentage inhibition was calculated after correction with cell-control and virus-control wells as follows: Inhibition (%) = [1 − (mean fluorescent foci in sample wells − mean fluorescent foci in cell-control wells) / (mean fluorescent foci in virus-control wells − mean fluorescent foci in cell-control wells)] × 100. The dilution corresponding to 50% inhibition was determined using the Reed–Muench method. The distance ratio was calculated as: d = (% inhibition above 50% − 50) / (% inhibition above 50% − % inhibition below 50%). The logarithm of the 50% inhibitory dilution was then calculated as: log_10_ (50% inhibitory dilution) = log_10_ (dilution above 50% inhibition) + d × log_10_ (dilution factor). IC_50_ values were calculated by dividing the starting antibody concentration by the 50% inhibitory dilution.

### Sequence analysis of RSV strains

Viral RNA extracted from RSV-positive clinical samples collected at Chongqing Medical University was reverse-transcribed into complementary DNA (cDNA). Whole-genome amplification was performed using a tiled amplicon strategy with overlapping primer pools and a high-fidelity DNA polymerase, generating overlapping fragments spanning the entire viral genome. Amplicons were subsequently subjected to next-generation sequencing (NGS; Illumina NextSeq 550) to obtain full-length genomic sequences for downstream analyses.

Raw sequencing reads (FASTQ) were processed using Trimmomatic (v0.39)^69^ for quality trimming, and read quality was assessed using FastQC (v0.12.1)^70^. Filtered reads were aligned to the RSV reference genome (RSV-A: EPI_ISL_412866; RSV-B: EPI_ISL_1653999, from GISAID^71^) using BWA (v0.7.17)^72^. Alignments were converted to BAM format, sorted, and indexed using SAMtools (v1.21)^73^, and alignment statistics and sequencing depth were calculated. Variant calling was performed using bcftools (v1.21; mpileup and call)^74^. A preliminary consensus sequence was generated based on detected variants for validation, and the final consensus genome used for downstream analyses was derived from the alignments using the SAMtools consensus function. A total of 1,687 RSV genomes were generated in this study spanning 2009–2025, including 941 RSV-A and 746 RSV-B isolates.

Publicly available sequences were retrieved from GISAID and NCBI. From GISAID (accessed on March 2, 2026), 50,965 RSV-A and 50,965 RSV-B sequences were obtained. From NCBI, 27,032 RSV-A and 22,448 RSV-B sequences were downloaded. Redundant sequences across the two databases were removed using SeqKit (v2.3.0)^75^, yielding 54,804 nonredundant RSV-A sequences and 39,675 RSV-B sequences. Fusion (F) protein sequences were extracted using BLAST+ (v2.13.0)^76^ based on reference sequences. This yielded a global dataset of 48,187 F protein sequences, including 27,013 RSV-A and 21,174 RSV-B sequences, incorporating those generated in this study using NGS.

For F protein analyses, sequences were aligned using MAFFT (v7.525)^77^, and mutation frequencies were calculated using custom shell scripts. For phylogenetic analysis of the F protein, sequences were aligned with MAFFT (v7.525), and duplicate sequences were further removed using seqkit. Maximum-likelihood phylogenetic trees were inferred using IQ-TREE (v2.2.6)^78^ under the GTR+G4+I substitution model. Branch support was assessed with 1,000 ultrafast bootstrap replicates and 1,000 SH-aLRT tests. All analyses were performed using 16 CPU threads, and trees were exported in Newick format.

### RSV cytopathic effect (CPE) inhibition neutralization assay of clinical isolates

RSV neutralization against clinical isolates was measured using a cytopathic effect (CPE) inhibition assay. Monoclonal antibodies were serially diluted twofold in maintenance medium and mixed with an equal volume of virus containing 100 CCID_50_ per well. The virus–antibody mixtures were incubated for 2 h at 36.5 °C in 5% CO_2_, after which HEp-2 cells were added to each well of a 96-well plate at 1–2 × 10^5^ cells mL^-1^. Plates were incubated at 36.5 °C with 5% CO_2_ for 5–7 days. Wells showing clear RSV-induced CPE were scored as infected. Neutralizing activity was defined as the antibody concentration that inhibited CPE in 50% of replicate wells. Because this assay is based on endpoint scoring across twofold serial dilutions rather than continuous dose–response curve fitting, neutralization values are discrete dilution-derived estimates. Therefore, identical neutralization values should be interpreted as endpoint-equivalent titers rather than precise continuous IC_50_ estimates.

### Phylogenetic clustering of antibody sequences

Antibody sequences were first annotated for V(D)J gene usage using AssignGenes.py from the Change-O toolkit (v1.3.3)^79^, and clonal lineages were defined with DefineClones.py based on Hamming distance model (--model ham) with a distance threshold of 0.25 (--dist). Sequences assigned to the same clone were aligned using MAFFT (v7.525) to generate multiple sequence alignments^77^. The resulting alignments were then used to infer phylogenetic relationships with IQ-TREE (v2.2.2.6)^78^. Model selection was performed using ModelFinder Plus (MFP) with the “-m MFP” option. The inferred trees were further processed and visualized with ETE3^80^. Subsequent mechanistic analyses used a single, consistent cross-reactive light chain derived from antibody CNR2360.

### Cryo-EM sample preparation and data collection

Purified RSV F proteins (prefusion and postfusion) at 1.0 mg mL^-1^ were incubated with a 1.2-fold molar excess of antibody Fab in 20 mM Tris pH 8.0, 200 mM NaCl at 4℃ for 30 min. Immediately prior to grid preparation, 0.05% N-Dodecyl-β-D-maltoside (DDM) was added to the sample. A 4 μL aliquot of the mixture was applied to glow-discharged holey carbon-coated gold grids (C-flat, 300 mesh, 1.2/1.3). Grids were blotted for 5 s with zero blot force at 100% relative humidity and plunge-frozen in liquid ethane using a Vitrobot (FEI). Micrographs were collected with a microscope. Movies were recorded using a direct detector with a defocus range between -1.2 μm and -2.0 μm. Automated single particle data acquisition was carried out by SerialEM. Full data collection parameters are reported in Table S1.

### Cryo-EM data processing

For the RSV F–Fab complex, movie stacks were motion-corrected using patch motion correction, and the contrast transfer function (CTF) parameters were estimated using patch CTF estimation in cryoSPARC^81^. Particle picking was performed using template-based picking with a previously determined RSV prefusion F structure as the template. Picked particles were visually inspected, extracted, and Fourier cropped before multiple rounds of 2D classification were performed to remove poorly aligned or contaminant particles. The remaining particles were re-extracted with box sizes of 320, 300, or 256 pixels depending on dataset requirements. Subsequently, ab initio reconstruction followed by heterogeneous refinement was performed to separate different particle populations. The selected particle subsets were further refined using homogeneous refinement and non-uniform refinement in cryoSPARC to improve map quality. Once the resolution plateaued and no further improvement was achieved by particle selection, the best particle subsets were subjected to local and global CTF refinement, followed by a final round of non-uniform refinement to generate the final reconstruction.

### RSV F + antibody Fab complex model building and refinement

Initial models of the RSV F–Fab complexes were generated using AlphaFold3^82^. The models were docked into the cryo-EM density maps using UCSF ChimeraX^83^. Manual model adjustment and rebuilding were performed in Coot^84^. The models were subsequently refined using real-space refinement in Phenix^85^ with secondary structure and geometry restraints. Iterative rounds of manual correction in Coot and refinement in Phenix were carried out to optimize model geometry and agreement with the density map.

### Statistical Analysis

Most statistical analyses were performed using R (v4.1.0). Statistical significance for Figures 1B–1C; Fig 5D; Figures S1B, S1D and S1E was assessed using the unpaired Mann-Whitney test, with Bonferroni correction applied for multiple comparisons. For Figure 5A, statistical significance relative to RSV was evaluated using Fisher’s exact test. Half-maximal inhibitory concentrations (IC_50_) in neutralization assays were calculated by linear interpolation between the two antibody concentrations surrounding 50% neutralization.

## Data availability

Cryo-EM maps in this study have been deposited in the Electron Microscopy Data Bank with accession numbers EMD-69567 (CNR2175), EMD-81090 (CNR2358), EMD-81092 (CNR3125 and CNR3184), EMD-81094 (CNR2473), EMD-81096 (CNR3017), EMD-81098 (CNR2298), EMD-81100 (CNR2185) and EMD-81102 (4D7). Models have been deposited in the Protein Data Bank with identification codes 24JN (CNR2175), 27EG (CNR2358), 27EH (CNR3125 and CNR3184), 27EJ (CNR2473), 27EL (CNR3017), 27EN (CNR2298), 27EP (CNR2185), 27ER (4D7).

## Acknowledgements

We thank X. Huang, X. Li, B. Zhu and L. Chen for cryo-EM data collection at the Center for Biological Imaging (CBI), Institute of Biophysics. We thank Y. Chen, Z. Yang and B. Zhou for technical support with BLI experiments. We sincerely thank all pediatric healthcare workers and toddlers from the Children’s Hospital of Chongqing Medical University who contributed blood samples for this study. We also acknowledge Nanjing Vazyme Biotech Co., Ltd. for technical assistance and valuable support throughout this work.

## Author contributions

X.W., Y.C., E.L., and W.H. designed and supervised the study. X.W., J.D., and Y.L. wrote the manuscript. L.X. and X.X. provided support for isolation and sequencing of antibodies. Y.L. analyzed the overall difference between pediatricians and toddlers. J.D., S.L. and F.J. conducted a detailed analysis of deep mutational scanning (DMS) data. J.D., K.X. and S.L. analyzed the antibody repertoire between pediatricians and toddlers based on DMS. K.X., J.D., W.Y., M.M., and L.W. collected Cryo-EM data, solved atomic structures, and made visualizations for the data in this study. Y.H. and Z.L. provided essential support for the isolation and sequencing of clinical RSV strains. J.N. performed the neutralization of mAbs against RSV isolates. J.W. and J.D. analyzed the prevalence of RSV or hMPV strains. J.D., Y.L., and K.X. designed the clone analysis and validation. M.M., Y.Z., and R.F. purified the F proteins of RSV and hMPV. H.Z. collected the blood samples and clinical specimens. L.X. provided guidance on evaluating the functional of the mAbs in vivo. Y.Z. and C.Z. provided writing suggestions during manuscript preparation. All authors reviewed and approved the final manuscript.

## Figures S1–S6

**Figure S1.**
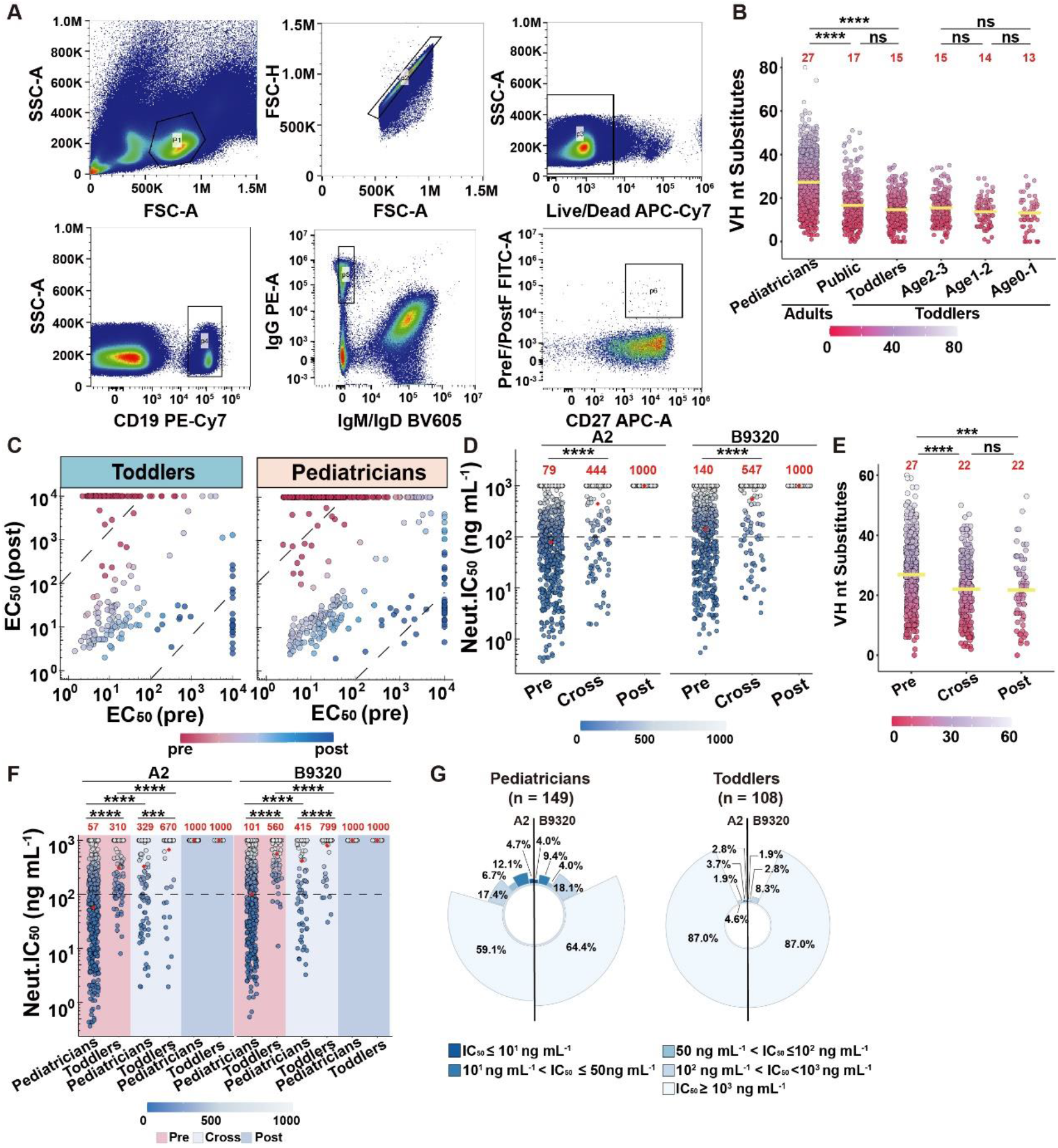
Age-associated maturation and functional profiling of RSV F-specific antibodies. **(A)** Gating strategy for a representative sample. Memory B cell reactivity with RSV F probes is shown in the lower right panel. SSC, side scatter area; FSC, forward scatter area. **(B)** Distribution of VH nucleotide substitutions in RSV F-specific memory B cells from adults and toddlers. Yellow lines and numbers above points indicate group means. **(C)** Scatter plot of binding affinities (EC_50_) of antibodies for preF (x-axis) and postF (y-axis). Each antibody is colored based on its EC_50_ (preF)/ EC_50_ (postF) ratio. Dashed diagonal lines indicate classification thresholds: antibodies above the upper line (≥100-fold preference for preF; EC_50_ (preF) ≤ 10⁴ ng mL^-1^ and EC_50_ (postF) ≥ 10^4^ ng mL^-1^) are defined as preF-specific, those below the lower line (≥100-fold preference for postF; EC_50_ (preF) ≥ 10^4^ ng mL^-1^ and EC_50_ (postF) ≤ 10⁴ ng mL^-1^) as postF-specific, and those between the lines as cross-reactive (0.01–100 ratio range) showing measurable binding to both preF and postF. **(D)** IC_50_ of these antibodies against RSV A2 and B9320. Red dots and the values above indicate the geometric mean IC_50_, the black dashed line marks the threshold for high neutralizing potency. **(E)** Distribution of VH nucleotide substitutions in preF-specific, cross-reactive and postF-specific antibodies; Statistical significance was determined by unpaired Mann-Whitney nonparametric test followed by Bonferroni correction for multiple comparisons. **(F)** Neutralization potency (IC_50_) of antibodies from adults and toddlers against RSV A2 and B9320 groupby binding preference. Red dots and values above indicate the geometric mean IC_50_ of antibodies with IC_50_ < 10^3^ ng mL^-1^, and the black dashed line marks the threshold for high neutralizing potency. **(G)** Proportions of cross-reactive antibodies against RSV A2 and B9320 in pediatricians and toddlers, grouped by neutralizing potency. ***p<0.001, ****p<0.0001, ns, no significance.

**Figure S2.**
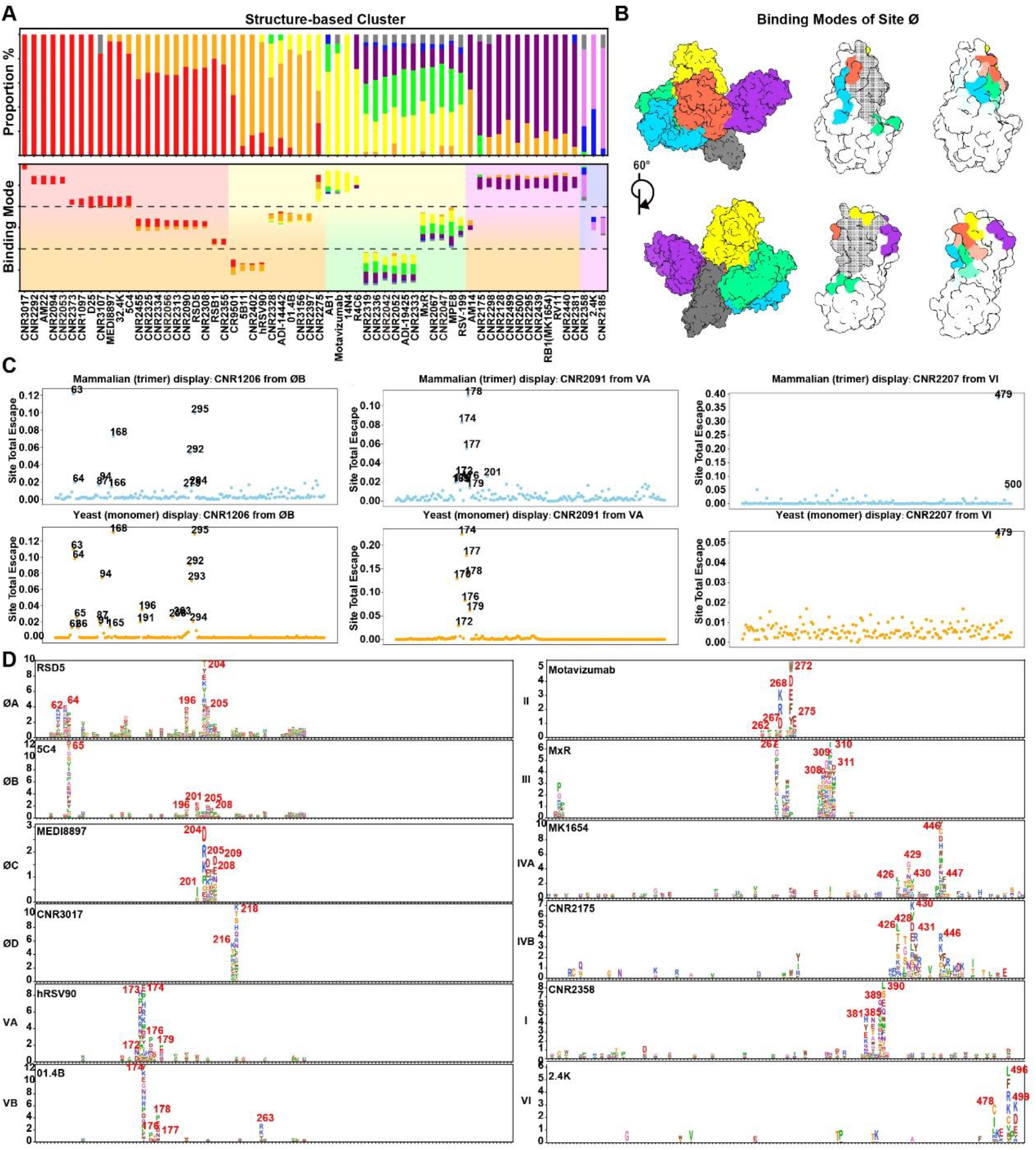
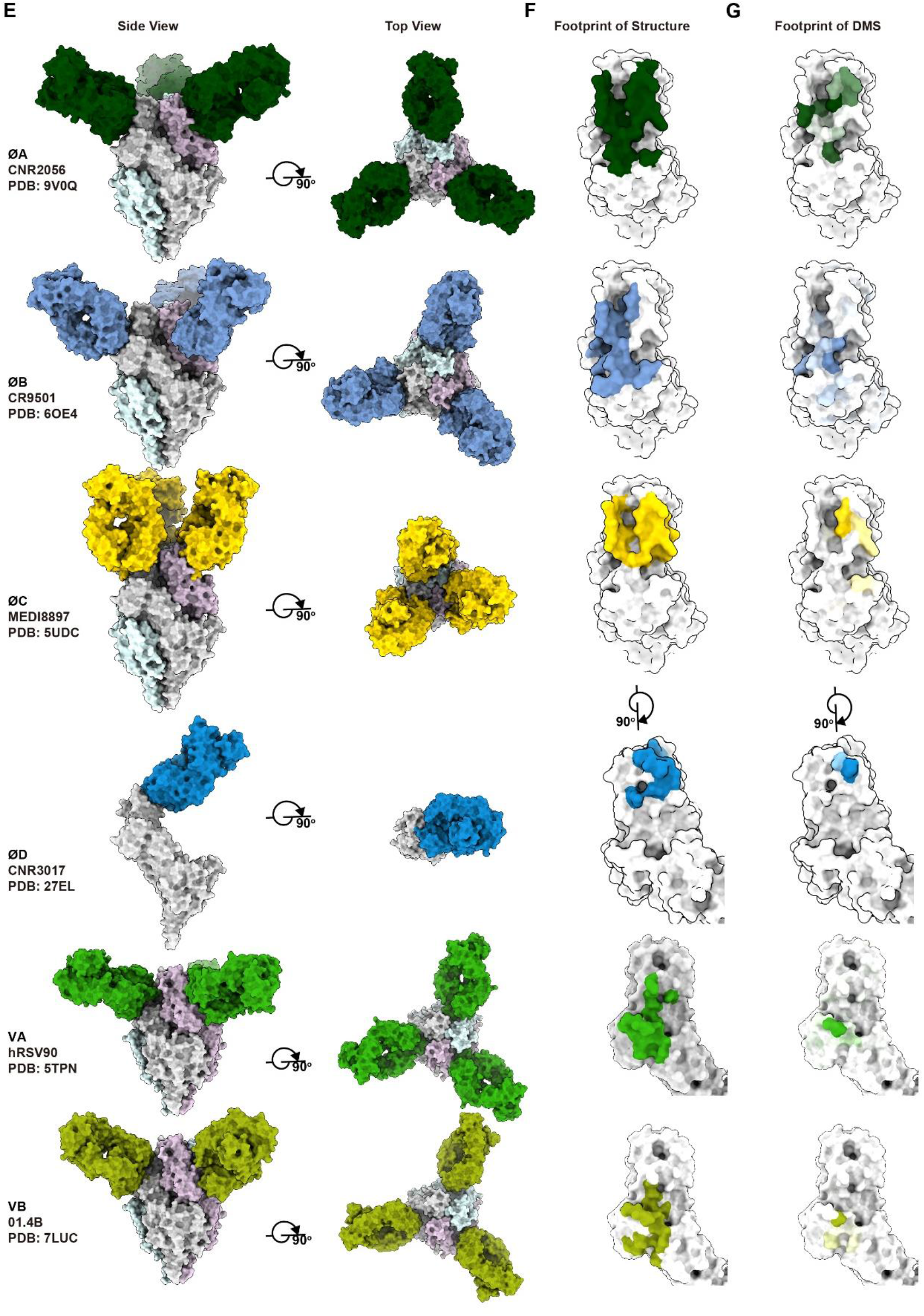
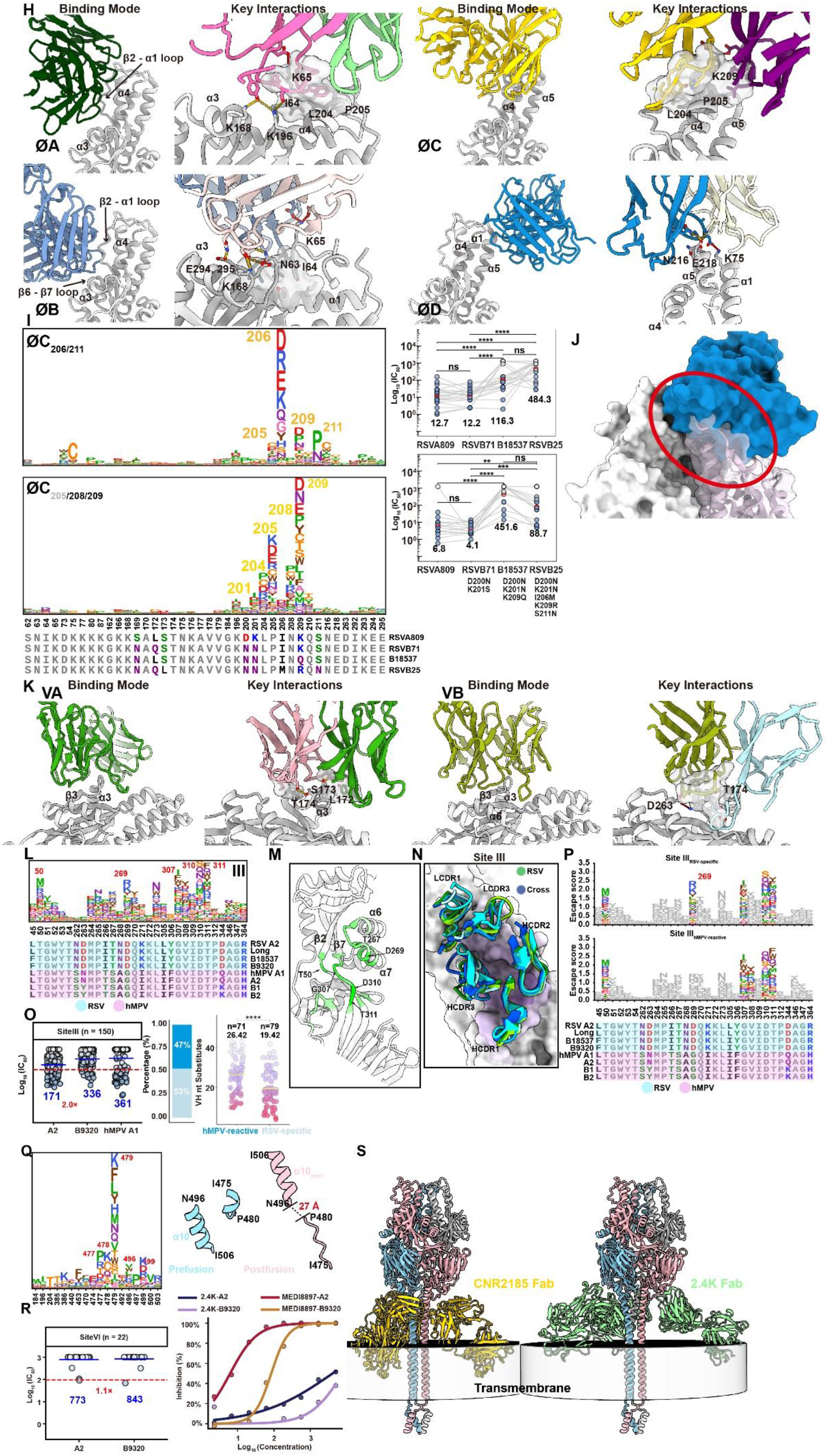
Integrated DMS and structural mapping defines the functional epitope landscape of RSV F. **(A)** Top: Stacked bar plot showing antibody clusters defined by structural data, sorted by classical epitopes (Ø: red, V: orange, II: yellow, III: green, IV: purple, I: pink, VI: grey). Each bar represents an antibody and indicates the distribution of its epitope residues across classical epitope regions. Bottom: Stacked bar plot showing the spatial positions of antibody epitopes. Bar length represents the number of epitope residues, colors indicate the classical epitope assignment of each residue. Bars are positioned according to structural overlap, binding modes, and functional differences among epitopes, such that antibodies with more similar binding modes are placed closer together. **(B)** Structures of five site Ø–targeting antibodies showing distinct binding modes and spatial overlap. **(C)** DMS point plots of representative antibodies showing concordance between yeast-displayed preF monomer and mammalian-displayed preF trimer platforms, including antibodies targeting sites Ø, V, and VI, which span from the head to the base of RSV F. **(D)** DMS escape logo plots for representative antibodies targeting each functional epitope. **(E)** Surface representations of representative antibody–RSV F complex structures for sites ØA, ØB, ØC, ØD,VA and VB, shown in side and top views. Fabs are colored by epitope-specific colors. **(F)** Structural footprints of representative antibody–RSV F complex structures. **(G)** DMS footprints of representative antibody–RSV F complex structures. **(H)** Binding modes and key interactions of antibodies targeting sites ØA, ØB, ØC, and ØD. Binding orientations are annotated with RSV F secondary structure elements, and key antibody–antigen interactions are labeled. **(I)** Escape logo plots and neutralization potency (IC_50_) of antibodies targeting the ØC subcluster against RSV clinical isolates harboring critical escape mutations, with RSV809 (subtype A) served as a control. Epitope amino acid multiple sequences alignment analysis for sites Ø (lower panel). Conserved residues are shown in gray. Variable residues are colored by physicochemical properties: hydrophobic residues (A, L, I, P, F, and M), black; polar residues (G, S, and T), yellow; neutral residues (Q and N), purple; basic residues (K, H and R), blue; and acidic residues (D and E), red. **(J)** Structural analysis of CNR3017 (PDB ID: 27EL) showing steric clashes between the Fab and the RSV F monomer. **(K)** Binding modes and key interactions of antibodies targeting sites VA (hRSV90, PDB ID: 5TPN) and VB (01.4B, PDB ID: 7LUC). Binding orientations are annotated with RSV F secondary structure elements, and key antibody–antigen interactions are labeled. **(L)** Escape mutation profiles and projection of RSV F binding activity derived from DMS for sites III, with key residues labeled. Epitope amino acid multiple sequences alignment analysis for sites III (lower panel). **(M)** Cartoon representation of RSV F colored by key DMS escape residues for site III, with representative residues and secondary structure elements labeled. **(N)** Binding convergence analysis based on published structural data for site III–targeting antibodies. Each antibody is shown in a distinct color. Site III antibodies are further distinguished by two color schemes representing hMPV-cross-reactive and RSV-specific antibodies. **(O)** Left: Neutralization potency (IC_50_) of site III–targeting antibodies against RSV A2, B9320, and hMPV A1. Middle: Proportion of hMPV cross-reactive antibodies among site III antibodies. Right: Comparison of heavy chain substitutions between hMPV cross-reactive and RSV-specific site III antibodies. **(P)** Escape mutation profiles of RSV F binding activity derived from DMS for site III, stratified by whether antibodies cross-neutralize hMPV A1. Logo plots show escape scores for each mutation, with only the top five escape sites highlighted. Epitope amino acid multiple sequences alignment analysis for sites III (lower panel). Conserved residues are shown in gray. Variable residues are colored by physicochemical properties: hydrophobic residues (A, L, I, P, F, and M), black; polar residues (G, S, and T), yellow; neutral residues (Q and N), purple; basic residues (K, H and R), blue; and acidic residues (D and E), red. **(Q)** Left: escape mutation profiles of RSV F binding activity derived from DMS for sites VI, with key residues labeled. Right: cartoon representation of site VI on RSV F in both pre-fusion (sky blue) and post-fusion (pink) conformations. **(R)** Left: Neutralization potency (IC_50_) of site VI antibodies against RSV A2 and B9320. Blue lines and values indicate geometric mean IC_50_ values of antibodies. Red dashed lines mark the threshold for high neutralizing potency (≤100 ng mL^-1^). Right: Dose–response regression curves for 2.4K and MEDI8897 against RSV A2 and B9320. **(S)** Cartoon representation of 2.4K (PDB ID: 8T7A) and CNR2185 (PDB ID: 27EP) in complex with full-length RSV preF, with the transmembrane region highlighted.

**Figure S3.**
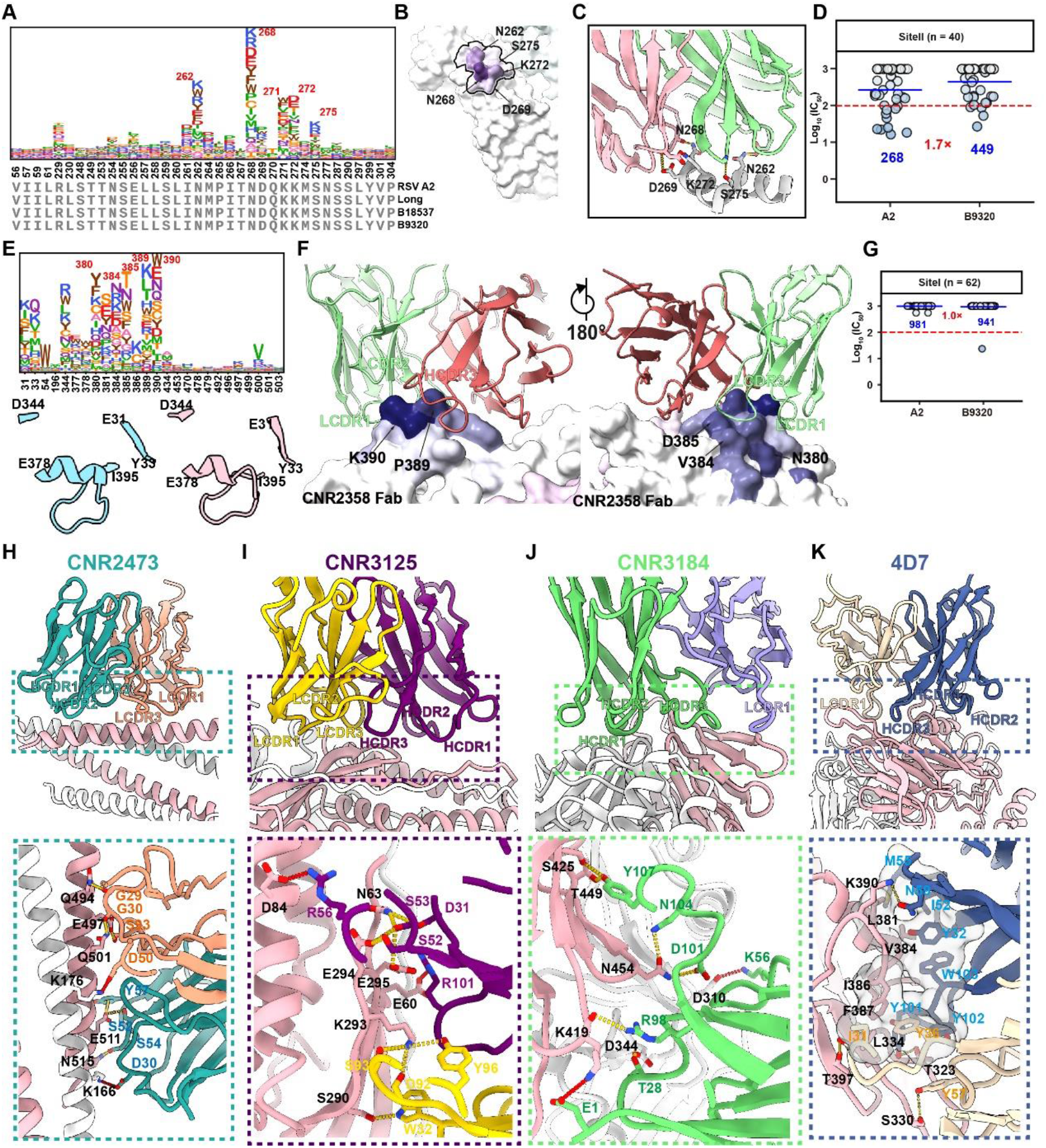
Cross-reactive and postF-specific antibodies reveal non-productive RSV F epitope landscapes. **(A)** Escape mutation profiles of RSV F binding activity derived from DMS for site II, with key residues labeled. Epitope amino acid multiple sequences alignment analysis for sites II (lower panel). Conserved residues are shown in gray. **(B)** Structural projection of RSV F binding activity derived from DMS for site II, with key residues labeled. **(C)** Binding mode of the site II antibody Motavizumab (PDB ID: 4ZYP), with key residues labeled. **(D)** Neutralization potency (IC_50_) of site II antibodies against RSV A2 and B9320. Blue lines and values indicate geometric mean IC_50_ values of antibodies. Red dashed lines mark the threshold for high neutralizing potency (≤ 100 ng mL^-1^). **(E)** Top: Escape mutation profiles of RSV F binding activity derived from DMS for site I with key residues labeled. Bottom: Cartoon representation of site I on RSV F in both pre-fusion (sky blue) and post-fusion (pink) conformations. **(F)** Binding mode of site I represented by antibody CNR2358 (PDB ID: 27EG). The antigen surface is colored by DMS escape scores for each residue, with critical residues labeled. **(G)** Neutralization potency (IC_50_) of site I antibodies against RSV A2 and B9320. Blue lines and values indicate geometric mean IC_50_ values of antibodies. Red dashed lines mark the threshold for high neutralizing potency (≤ 100 ng mL^-1^). **(H-K)** Detailed interactions between RSV post-F and antibody CNR2473 (PDB ID: 27EJ), CNR3125 (PDB ID: 27EH), CNR3184 (PDB ID: 27EH) and 4D7 (PDB ID: 27ER). Residues involved in hydrophobic patches and hydrogen bonds are shown as transparent surfaces and labeled, respectively. Heavy chain colors are consistent with those in Figure 3G.

**Figure S4.**
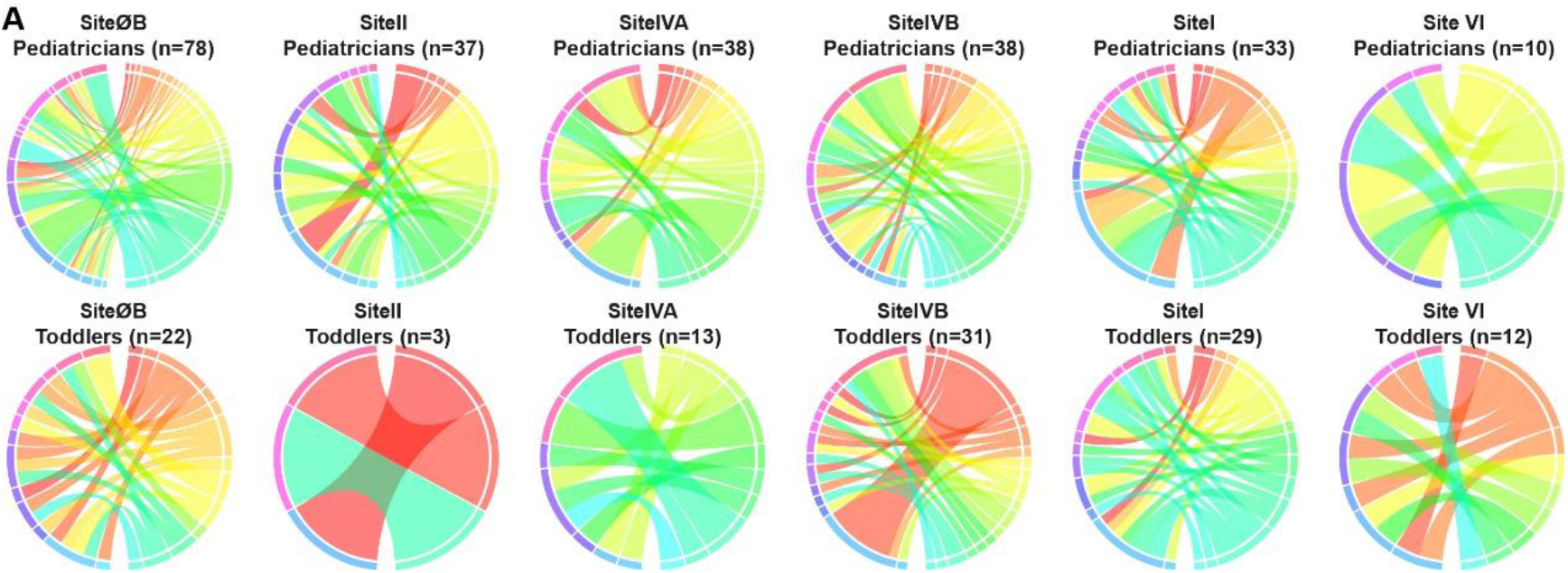
Germline pairing across RSV F antibody epitope groups. **(A)** Chord plot showing VH/VL germline pairing of sites ØB, II, IVA, IVB, I and VI antibodies in pediatrician and toddler groups.

**Figure S5.**
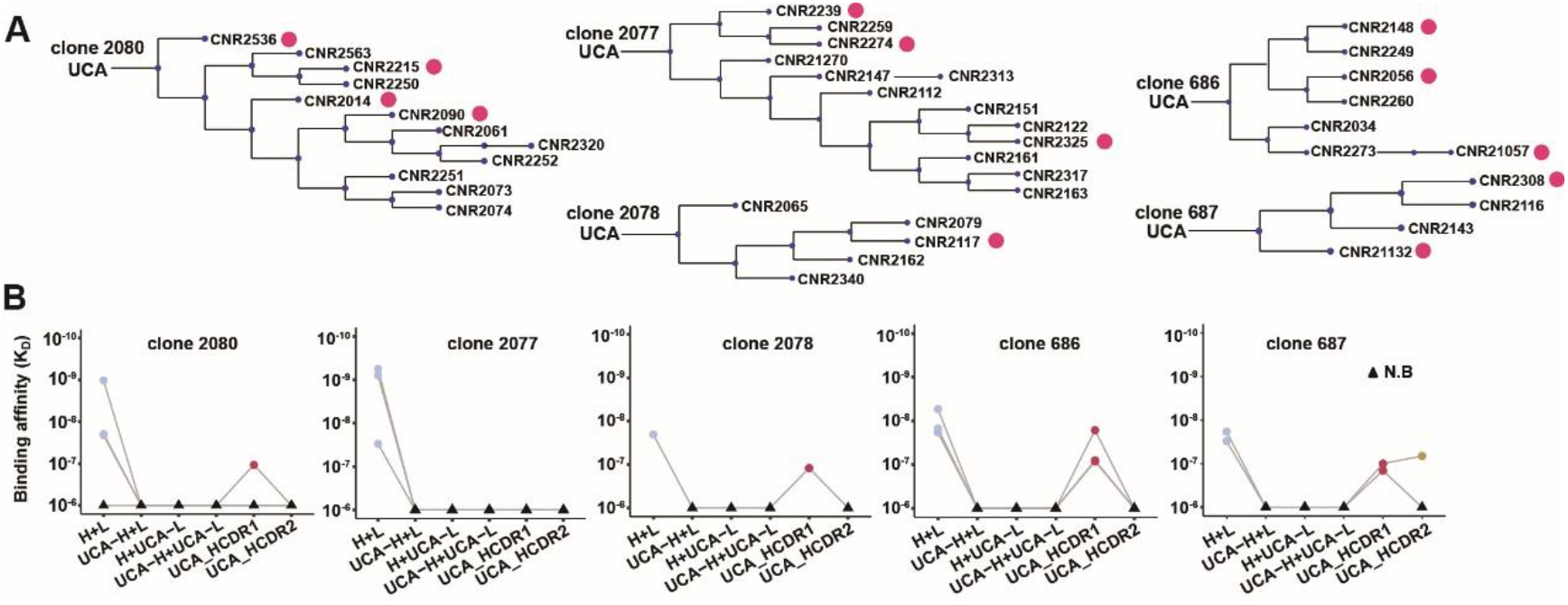
Germline reversion analysis of IGHV3-30/IGLV2-14 antibodies. **(A)** Genealogical trees of the IGHV3-30/IGLV2-14 antibody linage. Red dots indicate representative antibodies selected from each clone based on HCDR3 degeneracy for functional testing in Fig S16B. **(B)** RSV binding affinities of five antibody configurations derived from IGHV3-30/IGLV2-14 clones: H+L; UCA-H+L; H+UCA-L; UCA-H+UCA-L; the mature heavy chain containing UCA-reverted HCDR1 with the mature light chain (UCA-HCDR1); and the mature heavy chain containing UCA-reverted HCDR2 with the mature light chain (UCA-HCDR2). N.B., non-binding.

**Figure S6.**
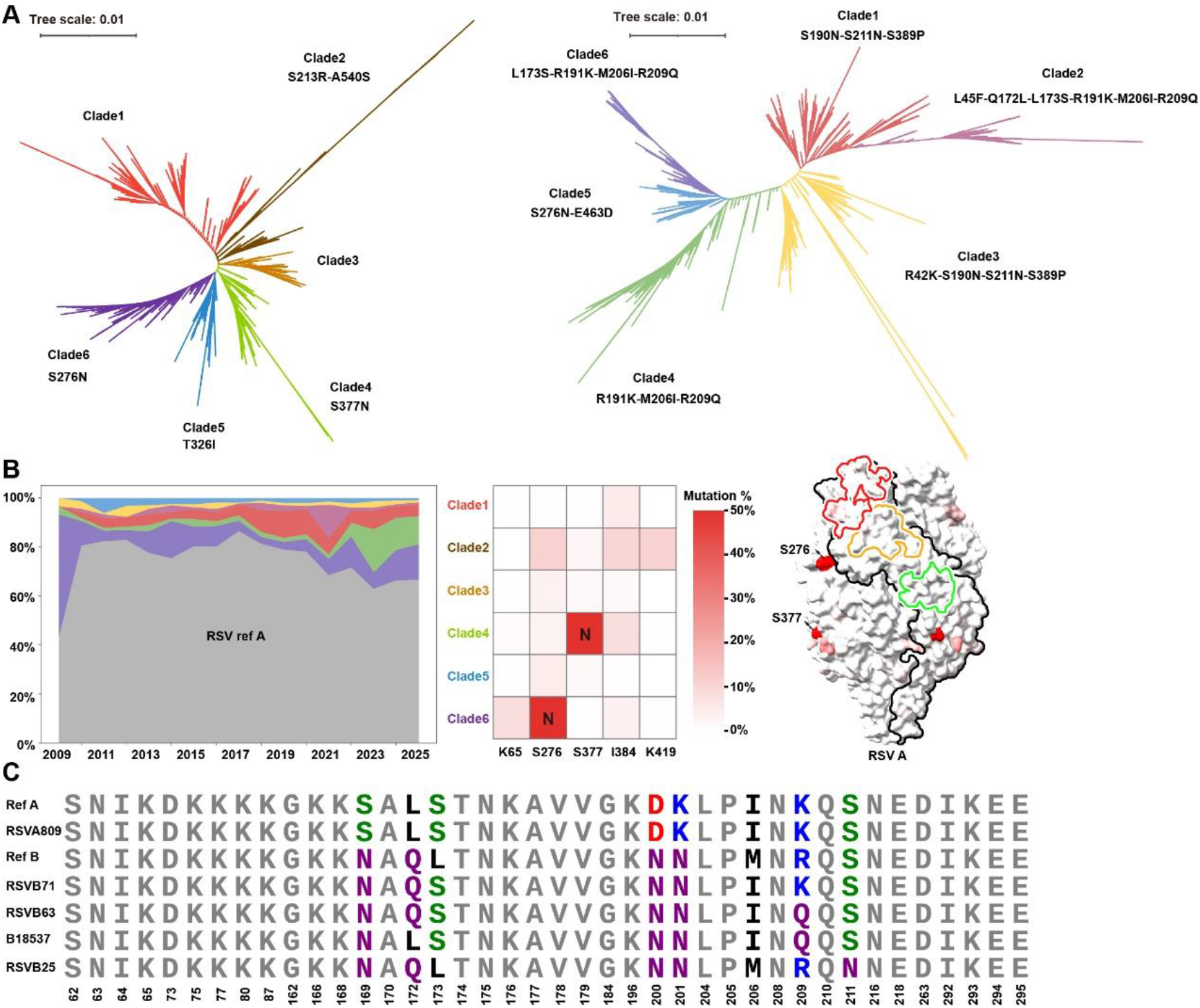
Clade dynamics and antigenic-site variation of RSV. **(A)** Phylogenetic tree and clade classification of RSV sequences. Maximum-likelihood phylogenies illustrating the genetic diversity of RSV-A (left) and RSV-B (right). Distinct clades (Clade 1–6) are color-coded, with representative signature mutations annotated for each clade. Scale bars indicate substitutions per site. Branch lengths reflect genetic distances, and nodes represent inferred ancestral relationships. **(B)** Temporal dynamics of RSV-A clades and mutation frequency across key sites. Stacked area plot showing the yearly distribution of RSV-A clades from 2009 to 2025. Each color represents a distinct clade, while grey denotes wild-type (WT) strains (left). Heatmap of mutation frequencies across major clades at key amino acid positions (65, 276, 377, 384, and 419). Color intensity reflects mutation frequency, with values exceeding 50% shown in red (middle). Surface representations of RSV F based on publicly available sequences of RSV-A. The color bar indicates mutation rate from 0% to 5%, with increasing red shading representing greater variability (right). **(C)** Multiple sequence alignment of representative RSV strains, including RSV-A and RSV-B reference strains, showing amino acid variation across epitope residues. Conserved residues are shown in gray. Variable residues are colored by physicochemical properties: hydrophobic residues (A, L, I, P, F, and M), black; polar residues (G, S, and T), yellow; neutral residues (Q and N), purple; basic residues (K, H and R), blue; and acidic residues (D and E), red.

**Table S1.** Cryo-EM data collection and atomic model refinement statistics, Related to Figures S2, S3 and Figure 3.

| Name | RSV pre-F monomer + CNR3017 + CNR3156 | RSV post-F + CNR2358 | RSV pre-F + CNR2298 | RSV pre-F + CNR2175 | RSV pre-F monomer + CNR2185 + CNR3107 + CNM3005 | RSV post-F + CNR247 3 | RSV post-F + CNR3125 + CNR3184 | RSV post-F + 4D7 |
| --- | --- | --- | --- | --- | --- | --- | --- | --- |
| **Data collection** |  |  |  |  |  |  |  |  |
| Microscope | Titan Krios 1 | Titan Krios G2 | Talos Arctica | Talos Arctica | Titan Krios 1 | Talos Arctica | Titan Krios 1 | Talos Arctica |
| Camera | K3 | K2 | K2 | K2 | K3 | K2 | K3 | K2 |
| Voltage (kV) | 300 | 300 | 200 | 200 | 300 | 200 | 300 | 200 |
| Total dose (e<sup>-</sup>/Å) | 60 | 60 | 60 | 60 | 60 | 60 | 60 | 60 |
| Symmetry imposed | C1 | C3 | C3 | C3 | C1 | C3 | C1 | C3 |
| **Data processing** |  |  |  |  |  |  |  |  |
| Micrographs (Total) | 3,740 | 2,081 | 2,939 | 2,696 | 4,619 | 2,812 | 2,884 | 2,438 |
| Micrographs (Used) | 3,654 | 1,851 | 2,676 | 2,326 | 4,474 | 2,812 | 2,796 | 2,164 |
| Particles selected | 2,201,814 | 1,985,525 | 1,011,638 | 934,278 | 4,488,385 | 4,068,813 | 3,771,136 | 1,427,338 |
| Particles included in final reconstruction | 140,616 | 323,542 | 49,863 | 29,429 | 187,010 | 709,983 | 116,164 | 575,229 |
| **Reconstruction** |  |  |  |  |  |  |  |  |
| Pixel size (Å) | 1.07 | 1.04 | 1.0 | 1.0 | 1.07 | 1.0 | 1.07 | 1.0 |
| Defocus range (μm) | -1.2 ~ -1.8 | -1.2 ~ -1.8 | -1.2 ~ -1.8 | -1.2 ~ -1.8 | -1.2 ~ -1.8 | -1.2 ~ -1.8 | -1.2 ~ -1.8 | -1.2 ~ -1.8 |
| Resolution (Å) (FSC=0.143) | 3.93 | 3.04 | 3.43 | 3.69 | 3.87 | 3.23 | 3.65 | 2.80 |
| **Model refinement** |  |  |  |  |  |  |  |  |
| Clash score | 9.80 | 7.83 | 10.79 | 15.00 | 8.77 | 14.52 | 6.75 | 13.43 |
| Rotamer outliers (%) | 0.00 | 0.00 | 0.00 | 0.00 | 0.00 | 0.00 | 0.00 | 0.00 |
| MolProbity score | 1.98 | 1.75 | 2.08 | 2.08 | 1.85 | 2.24 | 1.69 | 1.97 |
| **Ramachandran statistics** |  |  |  |  |  |  |  |  |
| Favored (%) | 92.48 | 95.31 | 90.58 | 94.08 | 94.52 | 89.20 | 95.48 | 95.26 |
| Allowed (%) | 7.52 | 4.69 | 9.42 | 5.92 | 5.48 | 10.80 | 4.52 | 4.74 |
| Outliers (%) | 0.00 | 0.00 | 0.00 | 0.00 | 0.00 | 0.00 | 0.00 | 0.00 |
| **R.m.s.deviation** |  |  |  |  |  |  |  |  |
| Bond lengths (Å) | 0.004 | 0.005 | 0.004 | 0.003 | 0.005 | 0.003 | 0.003 | 0.003 |
| Bond angles (°) | 0.653 | 0.539 | 0.622 | 0.642 | 0.763 | 0.609 | 0.562 | 0.570 |

